# A hidden protamine PTM code in sperm generates heterogeneous chromatin states and finetunes reproductive fitness

**DOI:** 10.1101/2025.11.18.688960

**Authors:** Lindsay Moritz, Christopher J. Woodilla, Ritvija Agrawal, Mashiat Rabbani, Samantha B. Schon, Wenxin Xie, Catherine A. Tower, Sowmya Srinivasan, Yi Sheng, Michael R. Baldwin, Patrick J. O’Brien, Rex A. Hess, Kyle E. Orwig, Sy Redding, Saher Sue Hammoud

## Abstract

Traditionally, the sperm genome is thought to be packaged by protamines into a uniformly compact and inert chromatin structure. Here, we challenge this long-standing view by demonstrating that protamine post-translational modifications (PTMs) present on distinct protamine molecules create discrete protamine-DNA chromatin states, ranging from weak to tightly associated chromatin configurations. Loss of these modifications alters protamine-DNA interactions *in vitro* and *in vivo*, compromising sperm chromatin integrity and impairing fertility. Therefore, these findings demonstrate that protamines do not merely serve as inert packaging proteins; rather protamine PTMs establish functional heterogeneity within sperm chromatin, creating compartment-like domains analogous to those in somatic cells. Thus, PTMs allow protamines to do more than simply compact the paternal genome—they likely encode a molecular blueprint that orchestrates the timely unpacking and reorganization of the paternal genome after fertilization.

## INTRODUCTION

Male fertility depends on the reliable production of sperm through spermatogenesis. This multi-phase process culminates in spermiogenesis, where developing spermatids undergo global cytoskeleton and chromatin remodeling to achieve a highly compact genome and competent sperm for fertilization.^1,2^ A central feature of this remodeling process is the replacement of canonical histones with testis-specific histone variants (testis-specific H2B (TH2B), TH2A, H3T, H2AL1/H2AL2, H1T2, and the spermatid-specific linker histone H1-like protein (HILS1))^3-9^, followed by incorporation of transition proteins and ultimately, protamines, which drive the final condensation of the sperm genome.^10-13^ Protamines are small, sperm-specific, highly basic nuclear proteins enriched in arginine residues, which mediate their non-specific binding to DNA.^14,15^ Most mammals express two forms of protamine: protamine 1 (P1) is expressed in its mature form, while protamine 2 is made as a precursor protein (pro-P2) that undergoes proteolytic processing to produce the mature form (P2) once bound to DNA.^16,17^ Importantly, both P1 and P2 are essential for proper sperm chromatin organization and male fertility; the loss of either protein^16,18-22^ or alterations in the P1:P2 ratio leads to decreased sperm motility and diminished fertilization rates.^5,16,23^

For decades, it was presumed that protamine-based chromatin packaging replaces most nucleosomes leading to a tight and uniformly packaged crystalline-like chromatin state consisting of repeating toroidal units of ≈50 kbp of DNA.^1,24,25^ This tight packaging presumably safeguards the paternal genome throughout its journey to the oocyte and erases any compartment-specific information that was engrained during germ cell development.^26,27^ In contrast to the nucleosome-based packaging in oocytes (and somatic cells), the protamine-condensed genome of sperm is largely cleared of parental chromatin marks, leaving only a small set of retained histones and persistent DNA methylation at loci critical for embryonic development and genomic imprinting.^28-31^ This differential packaging of oocytes and sperm raises a fundamental question: could the protamine-packaged regions also harbor a hidden layer of code or organization?

Protamine-based packaging is often considered a simple and passive molecular condensation program, relying on charge neutralization to hypercondense sperm DNA into a small nucleus.^1,14,32,33^ However, emerging data suggest that protamines engage in a regulated molecular compaction program with additional layers of encoded functional and structural information. For example, protamine post-translational modifications (PTMs) could influence local compaction patterns, higher-order folding, and, ultimately, regulate the accessibility and reprogramming potential of paternal chromatin following fertilization. Indeed, topological studies have shown that the sperm genome is organized into discrete ≈50 kbp loops, anchored to the nuclear matrix at specific sites known as matrix-associated regions, or MARs.^34-38^ Within these higher-order structures, it was shown that the coding region for the 5S rRNA clusters at the base of a loop, while satellite DNA is enriched within the loops themselves, suggesting a programmed and reproducible organization pattern.^38-41^ Despite these insights into the organizational framework, less is understood about the molecular mechanisms underlying this organization. Several early studies analyzing radiolabeled basic proteins from mouse and rat seminiferous tubules found that newly synthesized protamines are phosphorylated and many of these modifications appear to be removed during spermatid maturation, implying that dynamic protamine modifications play a regulatory role during spermiogenesis.^17^ A later study revealed that acquired phosphorylation of P1 in the early embryo is necessary for forming the male pronucleus, specifically promoting the exchange of protamine for histones.^42^ Recent biochemical and mass spectrometry analyses by our group and others demonstrated that protamines acquire specific PTMs during spermatogenesis that are stably retained in mature sperm.^43,44^ One such modification is acetylation of P1 at lysine 49 (K49ac), a rodent-specific modification whose loss impacts sperm function and fertility.^43^ Specifically, we have shown that substitution of K49 for alanine (K49A) alters DNA compaction and decompaction kinetics *in vitro*, and leads to premature male pronuclear decompaction and altered DNA replication kinetics in zygotes.^43^ Together, these data outline a regulatory role for protamine PTMs in both establishing the chromatin architecture in sperm and regulating protamine removal in the embryo. However, it remains to be determined how different PTMs coordinate to achieve this outcome and whether they give rise to distinct chromatin states in the sperm genome.

Here, using optical tweezers, we find that protamine works against low piconewton forces to compact DNA into regular structures *in vitro*. At higher forces, these structures relax in an all- or-none fashion, releasing discrete units of ≈30– 150bp of DNA. For P1 isolated from WT sperm, we observe two types of P1-DNA structures: a fast-relaxing, weak complex and a slower-relaxing complex, more resistant to mechanical disruption. Complementary salt fractionation experiments from testes and sperm reveal a similar dichotomy *in vivo*: a low-affinity, readily extractable P1-DNA population and a subset of high-affinity, salt-resistant P1-DNA complexes tightly bound to chromatin. Strikingly, these biophysical states are biochemically distinct *in vivo*: the low-affinity complexes are enriched for P1 K49ac, whereas the salt-resistant fraction is enriched for P1 phosphorylated at serine 43 and threonine 45 (S43/T45ph) and P2. Immunofluorescence analysis further reveal that these modifications appear to occupy spatially non-overlapping regions of the sperm nucleus, signaling that P1 modifications differentially compartmentalize chromatin within the protamine-bound genome. Functionally, substitutions at either K49^43^ or S43/T45 (new in this study) alter chromatin composition, condensation, and fertility. Interestingly, substitution of S43/T45 with non-phosphorylatable alanine residues (P1^S43AT45A^, AA) causes milder defects than phospho-mimetic, glutamic acid substitutions (P1^S43ET45E^, EE), which result in near-complete loss of sperm motility and infertility. These findings underscore that precise temporal control of P1 phosphorylation *in vivo* is essential for optimal fertility, but normal sperm production can still proceed in its absence. *In vitro*, both AA and EE proteins exhibit changes in the degree of condensation and mechanical stability of P1-DNA complexes. Specifically, alanine substitutions lead to more uniform, stable DNA packaging, while glutamic acid substitutions grossly impair DNA binding and packaging. Together, these data support a model in which protamine PTMs regulate the formation of biophysically and spatially distinct chromatin states during spermiogenesis. More broadly, our results challenge the longstanding view of sperm chromatin as uniformly organized within the nucleus and instead reveal an unappreciated layer of chromatin heterogeneity within the protamine-bound genome established by protamine modifications. This protamine-driven heterogeneity may not only create distinct packaging states within sperm but may also encode a temporal program for the stepwise unpacking of chromatin in the zygote, guiding the orderly transition of the genome from highly compacted in sperm to accessible in the embryo.

## RESULTS

### Protamine packages DNA in vitro into regular structures

Previously, we measured the rate at which mouse protamine compacts individual DNA molecules under laminar flow. We showed that compaction is a highly cooperative process where the rate and extent of compaction varies widely from molecule to molecule.^43^ This observation raised the question of whether the variability in compaction reflected underlying heterogeneity or complete lack of structural organization. To this end, we investigated P1-driven DNA organization through optical trapping experiments. In each experiment, we perform a series of three incubation-decompaction cycles (IDCs) (**Figure 1A**). In each IDC we first incubate a single DNA molecule in buffer alone or buffer containing P1 and hold it at an extension of 6μm for 3 minutes. Then, we initiate a force clamp at 8pN and measure the time-dependent decompaction of DNA (**Figure 1A**). The three IDCs are distinct; in the first IDC, both incubation and decompaction are performed in the presence of 150nM P1, in the second IDC, only incubation occurs with P1 in solution, and in the third IDC, both steps are performed entirely in the absence of P1 (**Figure 1A**). Generally, we compare the results from the first and third IDCs, as IDC2 is an intermediate case.

**Figure 1:**
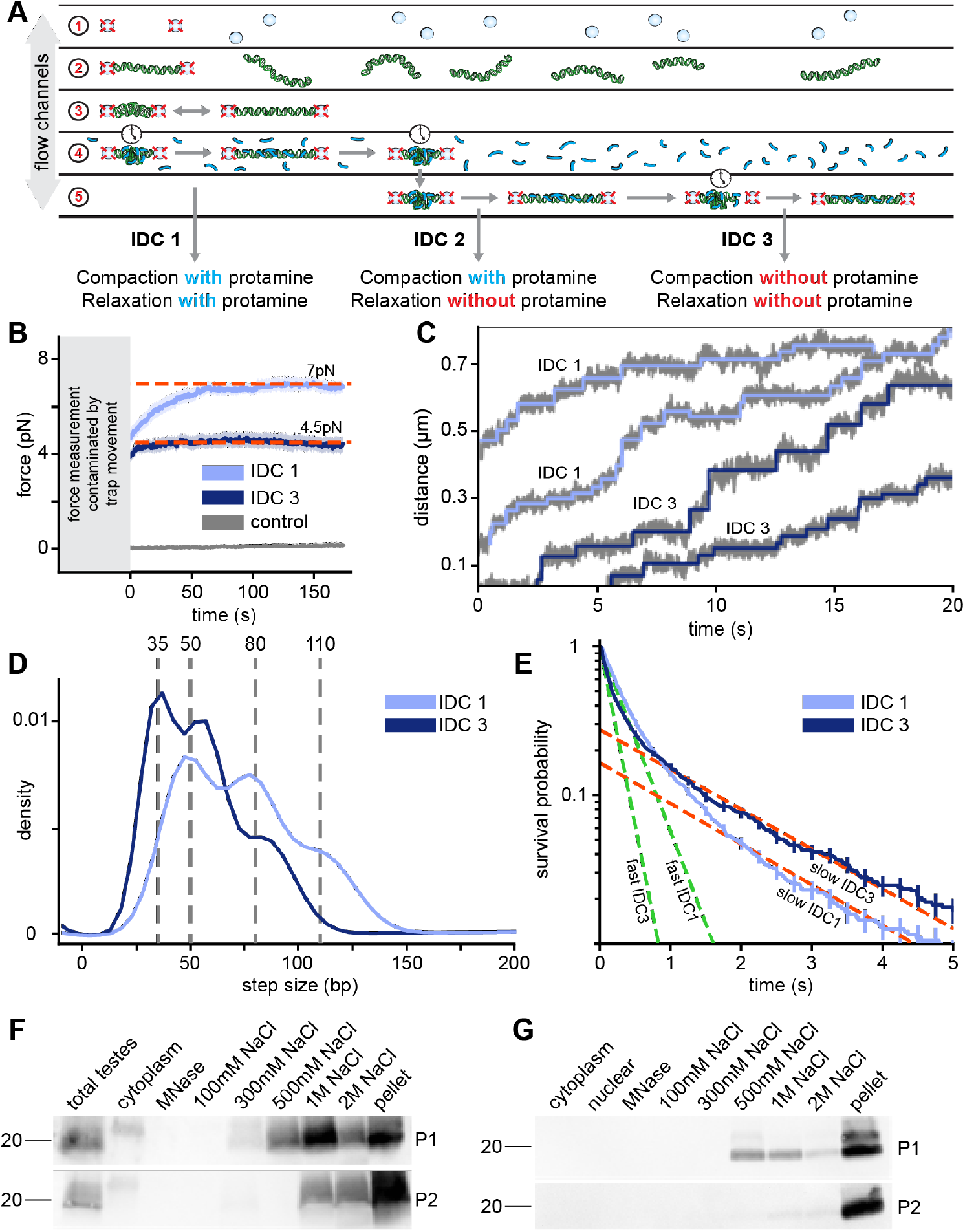
Protamine forms distinct but heterogeneous DNA complexes. **(A)** Schematic of optical tweezer experiments. Experimental procedure proceeds through the laminar flow channels in the following order (1) capture beads (2) capture DNA tether (move to buffer only channel) (3) control DNA IDC (move to protamine channel) (4) incubate in protamine and apply force clamp to measure decompaction, **IDC1**. Relax and incubate in protamine (move to buffer only channel) (5) apply force clamp to measure decompaction, **IDC2**. Relax and incubate in buffer and then apply force clamp to measure decompaction, **IDC3. (B)** Force creep measured during incubation phases of IDC1 and IDC3. Movement between flow channels and bead positioning prevents accurate measurement of the force leading to non-zero initial values (grey box). Red dashed lines indicate plateau force. Control sample contains no DNA. **(C)** Example traces showing P1-DNA decompaction steps from IDC1 and IDC3, data in grey and fitted steps in light (IDC1) and dark (IDC3) blue. **(D)** Joint histogram of P1-DNA decompaction steps from IDC1 and IDC3, grey dashed lines indicate peaks from Gaussian fits. **(E)** Survival probability of step dwell times for P1-DNA decompaction steps from IDC1 and IDC3. Dashed lines are derived from fit to bi-exponential decay. Green and red dashed lines indicate fast and slow decompaction step rates, respectively. **(F)** Western blots of P1 and P2 from salt fractionated P1-V5-tagged whole testes. **(G)** Western blots of P1-V5-tagged P1 and P2 from salt fractionated sperm.

During the incubation phase, and once the trap stabilizes, we measure changes in the force exerted on the DNA by protamine. Generally, we observe a gradual increase in force generated by the formation of P1-DNA complexes, a phenomenon we refer to as force creep (**Figure 1B**). We observe no force creep in controls lacking protamine (**Figure 1B**). In IDC1, the force creep stalls at ≈7pN, which sets the upper limit for protamine’s chromatin compaction power under these experimental conditions. Above this stall force, P1 may still bind to DNA, but is unable to compact DNA. We find that while the stall force is similar in the first two IDCs (in the presence of protamine), in the third IDC (in the absence of protamine), the force creep plateaus at ≈4.5pN (**Figure 1B**). This result reveals that a portion of protamine capable of recondensing the DNA remains bound even after a round of decompaction and extension in the absence of excess P1, illustrating that decompaction does not require dissociation.

In the decompaction phase, we increase the pulling force above the stall force to 8pN to promote decompaction. Once the force clamp stabilizes, we then measure time-dependent changes in the length of the DNA tether (**Figure 1C**). The time-extension traces reveal that protamine-DNA structures largely decompact in a stepwise fashion, releasing between 30 to 150bp per step (**Figure 1C,D**). Importantly, P1 appears to package DNA into distinct structural units (**Figure 1D, S1C,E**). In the first IDC, we observe peaks in the step size distribution at approximately 50bp, 80bp, and 110bp, which are all separated by, but are not integer multiples of 30bp (**Figure 1D**). In the third IDC, where incubation and decompaction are performed without P1 in the buffer, the likelihood of observing larger decompaction steps decreases, and while we still observe peaks at approximately 50bp and 80bp, a new peak emerges at 35bp (**Figure 1D**). Although decompaction is favored in our experiment due to the applied force (8pN) being close to the stall force (7pN), we still observe several “backward”, compaction steps (**Figure 1C, S1E**). While compaction steps are more variable in magnitude, their average peak size is comparable to that for decompaction steps, just in the opposite direction, reflecting similar structural principles govern both compaction and decompaction (**Figure S1E**). Together, these findings demonstrate that P1-DNA compaction and decompaction occur in a discrete, stepwise manner, organizing DNA into energetically preferred structural units. Since P1-DNA structures are regular, but not multiples of a clear “fundamental” packaging unit, this suggests that other factors, such as protamine composition or concentration in individual structures, competition for P1 binding sites, and DNA bending dynamics may also influence the size of P1-DNA structures.

Next, we examined the dwell time between each step. This is the amount of time that the system remains at a particular extension prior to a decompaction step. As evidenced by the non-linear decay of the natural logarithm of the step dwell survival distribution, there must be at least two rates of decompaction steps, suggesting the presence of at least two populations of compacted structures (**Figure 1E, S1D,F**). In IDC1 (performed with P1 in solution), we find a weak state, which persists for only ≈0.3s and stronger state that lasts on average for ≈1.5s, indicating that ΔΔG ≈ 1.5k_b_T more energy is required to unfold the stronger structures as compared to weaker structures (**Figure 1E, S1G**). Separating out these steps by size, either larger or smaller than 60bp, reveals that while, on average, smaller structures decompact faster than larger ones, both small and large structures exhibit both weak and strong states (**Figure S1H**). Moreover, compaction steps also display both slow and fast step dwell times and occur at rates similar to decompaction steps (**Figure S1I**). In IDC3 (performed without P1 in solution), the dwell time for the weaker state decreased roughly twofold, while the stronger state dwell time is largely unchanged (**Figure 1E, S1G**). This likely reflects partial dissociation of P1 from DNA and/or loss of cooperative activity of P1, consistent with the lower force plateau observed during IDC3 (**Figure 1B**). Interestingly, we observe an increase in the proportion of longer dwell times from IDC1 (19%) to IDC3 (28%) (**Figure S1G**). We suspect this is due to either weaker structures dissolving during the first IDC while stronger complexes persist to later IDCs and/or because weaker P1-DNA structures stabilize into stronger assemblies as the total incubation time increases.

Because each P1 is expected to bind ≈10bp of DNA^45^, the observed sizes of decompaction steps cannot be due to the release of a single P1 molecule, but likely arises from the cooperative relaxation of several P1 molecules simultaneously, or from a critical “linchpin” P1 molecule whose relaxation destabilizes the complete structure. The differences in dwell times we observe as a function of step size argues for cooperative action, assuming larger structures contain more P1 that all contribute an average energy to the overall structure. These observations raise an important question: do the distinct P1-DNA binding patterns we detect *in vitro* also occur *in vivo*, or is heterogeneity only a property of *in vitro* compaction of DNA by protamine? To this end, we examined if protamine creates multiple chromatin states *in vivo*. We performed subcellular and high salt fractionation in adult testes (**Figure 1F**) and sperm (**Figure 1G**) to assess the binding stability of protamine subpopulations. Immunoblot analysis of salt fractions reveals that P1 begins to elute at 300–500mM NaCl but progressively increases at higher salt fractions (**Figure 1F,G**). However, a subset of P1 remained in the insoluble pellet even after exposure to 2M NaCl (**Figure 1F,G**). In contrast, P2 is much more salt-resistant and largely remains in the insoluble pellet (**Figure 1F,G**). These distinct salt extraction profiles indicate that P1 and P2 can exist in chromatin states with different binding stabilities *in vivo*, both during spermatid chromatin remodeling and in mature sperm, consistent with our *in vitro* observations. Altogether, these results challenge the view that protamine packaging produces a uniform chromatin state and instead asserts that P1 establishes structurally regular but heterogeneous assemblies with distinct energies that can be potentially modulated by P2 incorporation and/or other mechanisms.

### Protamine-bound DNA fibers capture free DNA and enable interstrand sliding

Because P1 forms multiple regular sizes of compacted DNA structures, we surmised that during formation, nascent P1-DNA assemblies might migrate locally along the DNA to minimize DNA compaction energy and coalesce into larger assemblies. If true, we would also expect that any capture and sliding mechanism would also be able to operate across distal regions of the genome. To test this hypothesis, we performed experiments where we cycled between two laminar flow channels, one containing P1 and another containing fluorescently labeled 2.7kbp linear dsDNA molecules (**Figure 2A**). After first incubating with P1, we moved the DNA tether into the channel with fluorescent DNA where we readily observed long-lived foci corresponding to captured DNA (**Figure 2B-D**). When performing the same experiment in the absence of P1, we did not observe any fluorescent DNA capture (**Figure S2A**). Somewhat surprisingly, we observed that most of the captured DNA molecules were diffusive (horizontal line meandering) (**Figure 2C-E, S2C**). Quantification of these trajectories reveals that captured DNA molecules are able to slide for 100s of base pairs along the DNA tether at rates similar to those of proteins searching for DNA binding sites (**Figure 2E**)^46^. In some instances, we observed DNA switching between sliding and stationary states, suggesting that they had either coalesced with dark and stationary preexisting DNA structures or formed stable *de novo* structures (**Figure 2C, white arrow**). Moreover, we also observed collisions between and coalescence of fluorescent DNA foci (**Figure 2D, white arrows**). Together, these observations are consistent with an interstrand capture and sliding mechanism for the initial stages of sperm chromatin condensation.

**Figure 2:**
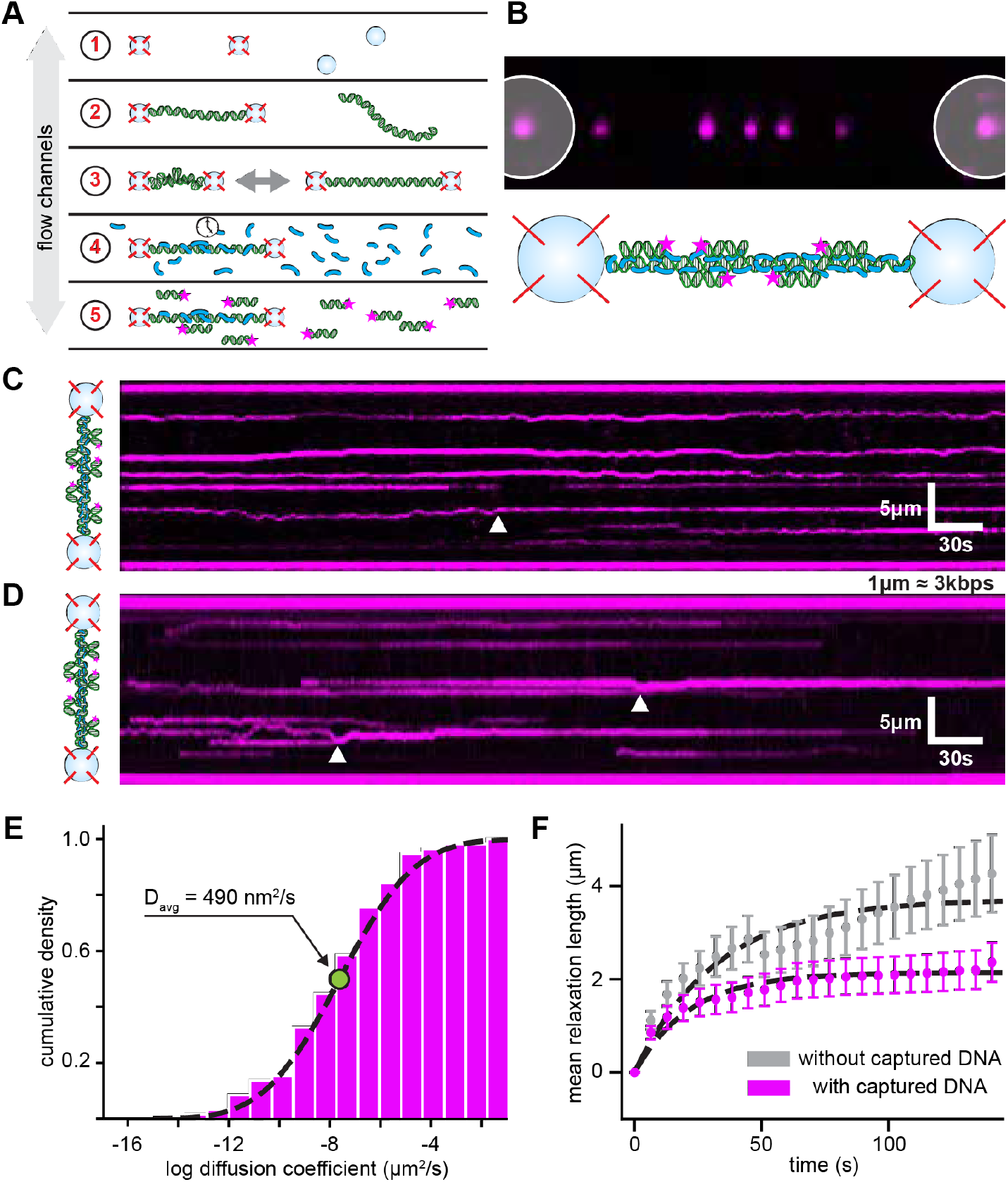
Inter-strand captured DNA is diffusive and stabilizes P1-DNA complexes. **(A)** Schematic of P1-DNA capture experiments. Details of experimental procedures are described in methods. **(B)** Example image (top) showing fluorescent captured DNA (magenta foci), scale bar is 3μm. The underlying DNA tether is unlabeled, and beads are indicated by white circles. (Bottom) Cartoon of captured DNA. **(C)** Example kymogram showing diffusion of captured DNA. A diffusion to capture event is indicated by the white arrow. **(D)** Example kymogram showing diffusion of captured DNA. Captured DNA coalescence indicated by white arrows. **(E)** Cumulative distribution of the natural logarithm of measured diffusion coefficients from tracked captured DNA. Arrow and value indicate the average diffusion coefficient. **(F)** Mean decompaction curves of DNA tethers in either buffer alone (grey) or 2.7kbp dsDNA (magenta), detailed experimental procedure described in methods.

The observation that captured DNA foci collide and coalesce implies a mechanism by which interstrand bridges might stabilize higher-order assemblies analogous to the manner in which larger structures exhibit longer step dwell times. As a result, we decided to test whether interstrand captured DNA could strengthen P1-DNA structures. We performed two IDCs, cycling between flow channels containing P1 or free unlabeled 2.7kbp linear dsDNA (**Figure 2A**). In the initial IDC (with P1 in solution), captured DNA adsorbed onto preformed P1-DNA structures, but was insufficient to statistically change the decompaction profile (**Figure S2B**). However, in the subsequent IDC (without P1 in solution), we observed a DNA capture-dependent reduction in the average decompaction rate (**Figure 2F**). Similar to the observation that excess P1 strengthens P1-DNA structures, we show that interstrand captured DNA can also reinforce P1-DNA structures when P1 is not readily available. Collectively, these data demonstrate that protamine-bound DNA, either locally during the initial steps of condensation or by capture of distal regions of the genome, creates weak structures that can slide along the captured interface until they are stabilized by either collision with preexisting P1-DNA structures or by additional loading of protamine.

### Protamine PTMs are established in a stage specific manner, enrich on different protamine molecules, and contribute to heterogeneous chromatin states

We and others had previously identified several PTMs on the C-terminus of P1.^43,47^ One intriguing possibility is that P1 molecules with distinct PTM states could explain the biochemically heterogeneous P1-DNA structures we observed both *in vitro* and *in vivo*. However, it remains unclear whether and how these modifications can generate distinct functional states or influence sperm chromatin architecture. Notably, we do know that loss of specific modifications, such as K49ac, disrupts chromatin structure and impairs reproductive fitness.^43^ However, in addition to K49ac, P1 is phosphorylated at S43 and T45 (**Figure 3A**). These two residues are conserved within the rodent lineage but are either absent or replaced by alternative residues in more distantly related species.^43^ To confirm the presence and dynamics of P1 S43/T45ph, we generated a polyclonal antibody against P1 S43/T45ph. To verify the specificity of the antibody and assess the function of these modifications, we generated two mouse lines: one with serine and threonine residues substituted with alanine (P1^S43AT45A^, referred to hereafter as AA) to model the loss of phosphorylation, and another with glutamic acid substitutions (P1^S43ET45E^, referred to as EE) to mimic ectopic or premature phosphorylation and to distinguish between the effects of phosphorylation and charge state (**Figure 4A**). In both models, we confirmed the successful introduction of the target mutations and verified the absence of any off-target genetic alterations (**Figure 4A, S4A,B**). In immunoblots of protein lysates from testes, we found that our antibody detected a clear band that was absent in protein lysates from AA and EE males (**Figure S3A**). Moreover, this band was outcompeted by a phosphorylated P1 peptide, but not an unmodified P1 peptide or a non-specific peptide, further confirming specificity of our antibody (**Figure S3B**).

**Figure 3:**
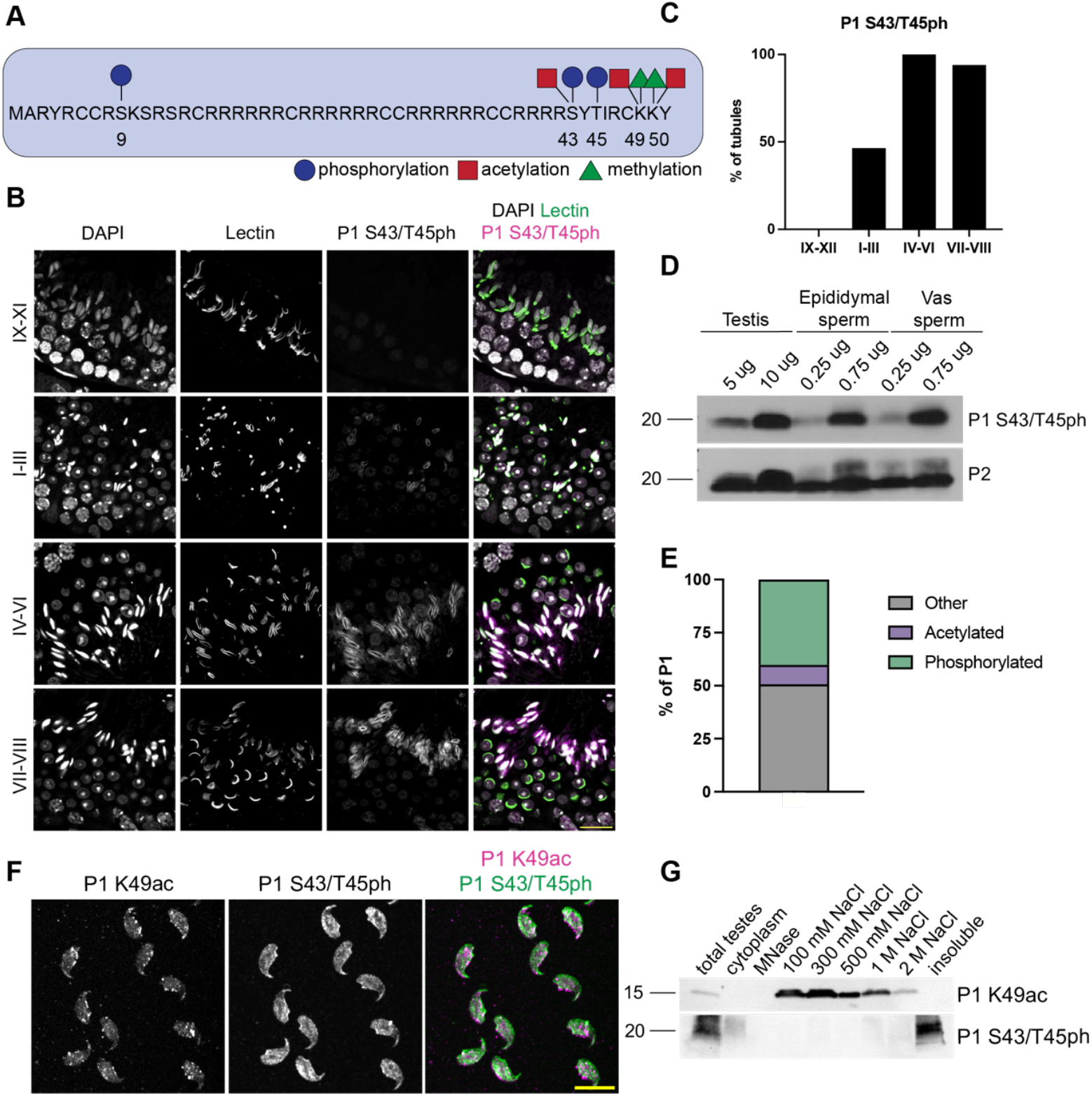
P1 modifications are established in a stage-specific manner and allow for distinct chromatin states. **(A)** Schematic representation of post-translational modifications identified by bottom-up mass spectrometry on mature mouse sperm. **(B)** Immunofluorescence staining of P1 S43/T45ph in adult testes cross sections using PNA-Lectin as a marker of the acrosome. Representative images from n=5 mice per timepoint. Scale bar: 20 μm. **(C)** Quantification of P1 S43/T45ph stage specificity across the seminiferous epithelial cycle illustrates establishment beginning in I-III, peaking in IV-VI and persisting into VII-VIII. A total of n=246 tubules were counted across all stages from n=5 mice. **(D)** Immunoblot of protein lysates from adult testes, epididymal sperm, and sperm from the vas deferens highlights persistence of P1 S43/T45ph from the testis to mature sperm. Depicted is a representative blot from n=2 independent experiments. **(E)** Percent of total P1 peptides identified by bottom-up mass spectrometry that were acetylated, phosphorylated, or either methylated or unmodified. Depicted data is from n=2 independent experiments and for each independent experiment, n=3 male mice were used, and sperm was pooled together for each. **(F)** Immunofluorescence of mature sperm stained for P1 K49ac and P1 S43/T45ph. Shown are representative images from n=3 independent experiments. Scale bar: 10 μm. **(G)** Immunoblot of protein lysates from subcellular and high salt fractionated mature sperm probed for P1 K49ac or P1 S43/T45ph. Shown is a representative blot with similar results obtained from n=4 independent experiments.

**Figure 4:**
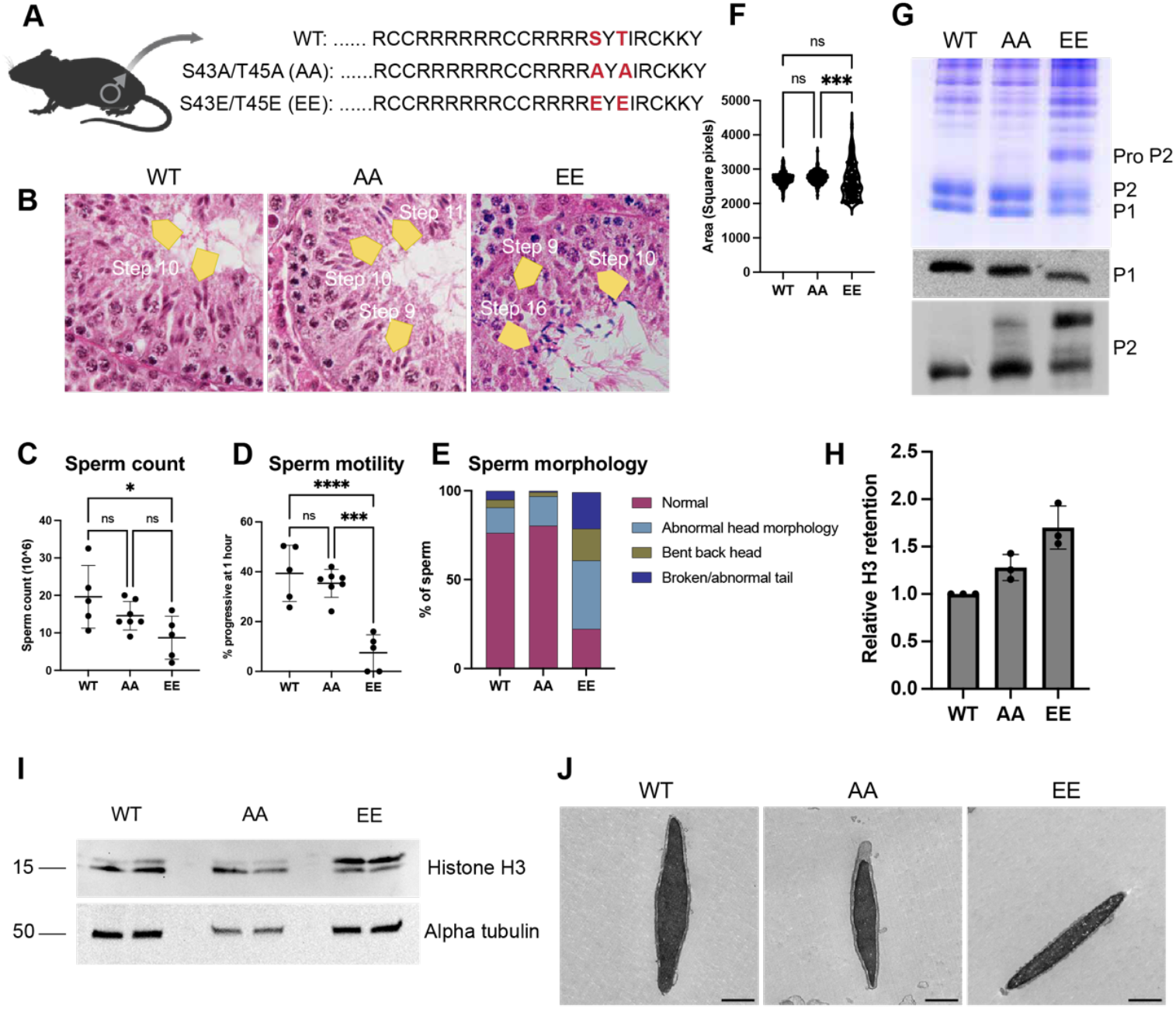
Loss of P1 S43/T45 phosphorylation compromises spermatid maturation, chromatin packaging, and sperm parameters. **(A)** Schematic of mutations made to introduce non-modifiable (AA) or phosphomimetic residues (EE) in the C-terminus of mouse P1. **(B)** Hematoxylin and eosin-stained adult testes from P1^+/+^ (WT), P1^S43AT45A^ (AA), and P1^S43ET45E^ (EE) males highlights defects in spermiogenesis present in both mutants. Yellow arrows indicate specific spermatid populations, highlighting a mixing of stages in both mutants and a failure of spermiation in EE males. **(C)** Total epididymal sperm count from WT (n=5), AA (n=7), and EE (n=5) males. Each dot represents the measurement of a single animal. Statistical tests were performed using a one-way analysis of variance (ANOVA) and adjusted for multiple comparisons, p=0.03 between WT and EE. Center line represents the mean and error bars represent standard deviation. **(D)** Progressive sperm motility from WT (n=5), AA (n=7), and EE (n=5) males. Each dot represents the measurement of a single animal. Statistical tests were performed using ANOVA and adjusted for multiple comparisons, p=0.0038 between WT and EE and p=0.000015 between AA and EE. Center line represents the mean and error bars represent standard deviation. **(E)** Quantification of abnormalities observed in WT, AA, and EE sperm. **(F)** Overall area of sperm across genotypes. A total of n=358 sperm from n=3 WT males, n=227 sperm from n=3 AA males, and n=174 sperm from n=3 EE males were counted. **(G)** Coomassie-stained acid urea gel (top) and immunoblots for P1 and P2 (bottom) highlight a defect in P2 processing in both mutants. **(H)** Quantification of relative H3 retention in both mutants compared to WT sperm. Each dot represents the average of n=3 technical replicates for a single animal. A total of n=3 males per genotype were used for analysis. **(I)** Representative immunoblot showing increased histone retention in both mutants. **(J)** Representative transmission electron microscopy images of WT, AA, and EE sperm. Scale bar: 1 μm

To define the stage or stages of the seminiferous epithelial cycle in which S43/T45ph is established, we co-stained adult testes cross-sections using our custom antibody with the acrosomal marker PNA-Lectin. Unlike K49ac, which is established in stages IX-XI of the seminiferous tubule cycle (**Figure S3C**), S43/T45ph is initiated in stages I-III (mid-stage spermatids) and peaked at stages IV-VI (late-stage spermatids, 100% of tubules). Both P1 K49ac^43^ and S43/T45ph persist in mature sperm (**Figure 3B-D**). This P1 S43/T45ph staining pattern was detected only in WT testes and was absent in both the phosphorylation-deficient (AA) and phosphomimetic (EE) mutant backgrounds (**Figure S3D**). In short, the differences in PTM establishment timing suggest a distinct temporal regulatory role in chromatin remodeling. Moreover, by generating and validating phospho-mutant mouse lines and a specific antibody against P1 S43/T45ph, we can begin to interrogate how these modifications regulate sperm chromatin architecture and male fertility.

Since these P1 modifications are established at different steps of spermatid maturation, we next examined whether these modifications occur on the same peptide or are enriched on distinct subsets of P1 molecules. To address this, we reanalyzed our previously published mass spectrometry data and found that P1 S43/T45ph is among the most abundant modifications, accounting for ≈40% of protamine peptides, whereas acetylation is present on ≈10%, in agreement with previous studies (**Figure 3E**).^47^ Notably, phosphorylation and acetylation are observed on different peptides, demonstrating for the first time that the protamine pool is compositionally heterogeneous with different populations manifesting distinct PTMs. Consistent with this observation, sequential high-resolution imaging reveals distinct nuclear localization patterns for these two modifications in mature sperm: S43/T45ph is enriched at the nuclear periphery, while K49ac is more broadly distributed in a punctate pattern throughout the nucleus (**Figure 3F**). We next asked whether these modifications confer differential DNA-binding affinities *in vivo*. Immunoblot analysis of high-salt fractions from testes (**Figure 3G**) and sperm (**Figure S3E**) revealed that P1 S43/T45ph remains associated with chromatin even after a 2M NaCl wash, whereas K49ac dissociates at much lower salt concentrations and is largely absent from the insoluble pellet (**Figure 3G, S3E**). These results show that differentially modified P1 molecules confer markedly different DNA-binding stabilities and chromatin states *in vivo* that may also differentially combine to produce the weak and strong binding states observed *in vitro*. Although both acetylation and phosphorylation reduce the positive charge on P1, which would be expected to weaken P1-DNA interactions, phosphorylated P1 is remarkably stable on chromatin. To explore potential mechanisms for this stabilization, we performed immunoprecipitation assays and found that P1 S43/T45ph, but not K49ac, interacts with P2 (**Figure S3F**). Furthermore, S43/T45ph shows enriched colocalization with P2 in mature sperm (**Figure S3G**). Together, these findings reveal that P1 modifications occur on separate P1 molecules, generating a heterogeneous protamine pool with different chromatin-binding stabilities and spatial distributions, and dictate interactions with P2 to stabilize chromatin binding.

### Loss of P1 S43/T45 phosphorylation compromises spermatid maturation

Next, to assess how P1 S43/T45ph influences sperm development and function, we analyzed AA and EE mutant mice (**Figure 4A and S4A,B**). While the testes-to-body weight ratios were comparable across genotypes (**Figure S4C**), histological examination of testis cross sections revealed significant delays in spermatid maturation in both mutants (**Figure 4B**). Typically, in wild-type testes, spermatid development is tightly synchronized within each tubule (**Figure 4B, left panel**). In contrast, AA and EE mutants had mixed populations of spermatid steps (e.g., steps 9, 10, and 11) within individual tubules (**Figure 4B**; middle and right), suggesting that the stage-specific acquisition of P1 S43/T45 phosphorylation is critical for maintaining an ordered and timely development of spermatids. Importantly, the defects observed in the testis cannot be attributed to differences in total P1 expression, as protein levels are similar across genotypes (**Figure S4D**).

Given the intratesticular defects, we next examined how S43/T45 substitutions impact mature sperm quality and morphology. Sperm analysis revealed a slight decrease in sperm count in AA males and a significant decrease in EE males (**Figure 4C**). Additionally, progressive motility is significantly reduced in EE males (**Figure 4D**), whereas the motility in AA sperm is comparable to WT (**Figure 4D**). Furthermore, morphological assessment of sperm reveals that ≈70% of EE sperm display head abnormalities and ≈20% display tail defects (**Figure 4E**). In contrast, the frequency of head defects in AA sperm is similar to WT (**Figure 4E**). Likewise, nuclear morphology assessment revealed notable differences between genotypes: AA sperm nuclei are uniform and similar in size and shape to WT sperm, whereas the EE sperm nuclei are heterogeneous—some nuclei are enlarged and decondensed while others are smaller with elongated heads (**Figure 4F**). Therefore, these observations suggest that while phosphorylation is not essential for sperm maturation, its absence in AA males leads to maturation delays that ultimately yield grossly normal sperm. In contrast, premature or ectopic phosphorylation, modeled in EE mice, leads to profound defects in sperm function, indicating that precise regulation of P1 S43/T45 phosphorylation is important for normal sperm morphogenesis and function.

### Substitution of P1 S43/T45 to either alanine or glutamic acid differentially affects sperm chromatin composition and compaction

Because sperm morphology is intimately linked to chromatin packaging, the structural abnormalities observed in EE sperm prompted us to examine whether chromatin remodeling and composition differ across genotypes. Histone-to-protamine exchange proceeds through a stepwise cascade, beginning with histone hyperacetylation and eviction, followed by incorporation of transition proteins (TNP1 and TNP2), and ultimately protamine deposition.^10,11^ To assess whether these intermediate steps were affected, we co-stained testis cross-sections for acetylated H4 and TNP2. Immunofluorescence analysis revealed similar levels and localization patterns of these markers across all genotypes, suggesting that initiation and progression of histone-to-protamine exchange proceed normally in both mutants (**Figure S4F**). We next examined the final chromatin composition in mature sperm by performing acid extraction followed by protein immunoblots. In AA mutant sperm, while the P1:mature P2 ratio was slightly reduced, the P1:total P2 ratio (accounting for pro-P2) was comparable to WT (≈1:2, **Figure 4G, S4E**). On the other hand, EE mutant sperm exhibit a slightly reduced P1:mature P2 ratio in addition to a P1:total P2 ratio greatly exceeding 1:2 (**Figure S4E**, ≈1:3), likely as a result of the accumulation of pro-P2 combined with a subtle reduction in P1 **(Figure 4G**). This decrease in P1 likely reflects reduced stability of the EE variant on chromatin, as total P1 protein abundance in testes was equivalent across genotypes (**Figure S4D**). Notably, both AA and EE sperm accumulated increasing amounts of pro-P2, indicating defective P2 processing **(Figure 4G)**. Furthermore, both mutants display incomplete histone eviction, as evidenced by elevated levels of retained histone H3 in mature sperm (**Figure 4H,I**). Consistent with these compositional changes, transmission electron microscopy revealed graded disruptions in chromatin packaging. WT sperm displayed uniformly electron-dense chromatin—characteristic of mature, fully condensed sperm nuclei (**Figure 4J, left**). In AA sperm, the chromatin remains largely condensed but shows uneven density, likely reflecting decompaction or delayed condensation (**Figure 4J; middle**). Strikingly, EE sperm showed highly heterogeneous and disorganized chromatin with alternating dense and lucent domains throughout the nucleus, indicative of aberrant chromatin architecture (**Figure 4J, right**). Interestingly, despite normal morphology, counts, and motility in AA sperm, the chromatin composition of these mutants displayed intermediate molecular and phenotypic features, falling between WT and EE sperm. Together, these findings highlight that precise temporal control of P1 phosphorylation is critical for robust, programmatic chromatin remodeling and spermatid maturation, with deviations leading to stochastic defects that compromise sperm chromatin integrity and structural fidelity.

### The substitution of P1 S43/T45 for glutamic acid, but not alanine, decreases DNA binding ability of P1

To better understand how P1 substitutions affect DNA binding behavior, either alone or in combination with P2 (both mature or unprocessed), we purified P1 and P2 from mature sperm from P1^+/+^ (WT P1), P1^S43AT45A^ (AA), and P1^S43ET45E^ (EE) mice (**Figure S5A**). Using electrophoretic mobility shift assays (EMSA; measuring binding to ≈300bp dsDNA) or fluorescence anisotropy (measuring binding to a 99bp fluorescent dsDNA), we show that AA has equivalent DNA binding affinity to WT P1 (**Figure 5A, S5B,C**). In contrast, EE displays weaker DNA binding affinity, evidenced by a dissociation constant (K_D_) of 1.1 µM compared to 0.43 µM for WT P1 and 0.61 µM for AA. This result demonstrates that the glutamic acid substitutions in the EE protein disrupt its ability to bind DNA effectively. Importantly, extending the incubation time in EMSA assays did not alter the binding kinetics, indicating that measured differences reflect changes to the intrinsic binding properties of protamine proteins at equilibrium (**Figure S5F**). We next investigated the DNA binding efficiency of P1 with mature P2 (purified from WT sperm) or unprocessed P2 (purified from EE sperm) by EMSA. Both WT P1 or AA with mature P2 bound DNA more efficiently together than either P1 protein alone (**Figure S5D, E**). However, WT P1 and AA showed slightly reduced binding efficiency when paired with pro P2 (**Figure S5D, E**). In contrast, EE bound with significantly lower efficiency in combination with either mature P2 or unprocessed P2 (**Figure S5D, E**).

**Figure 5:**
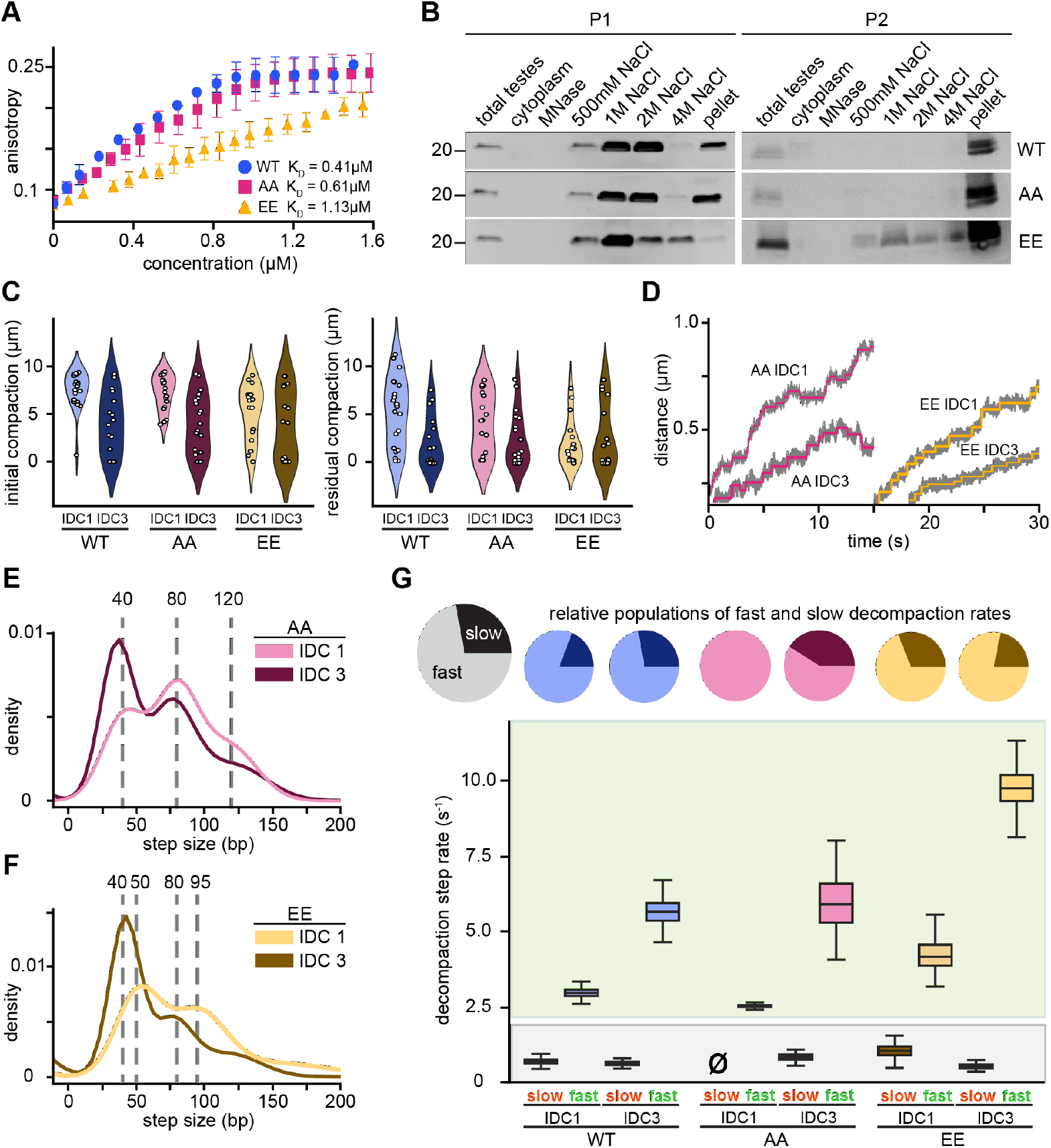
Substitution of P1 S43/T45 affects P1-DNA binding and dynamics. **(A)** Fluorescence anisotropy curves for WT P1, AA, and EE. **(B)** Western blots of P1 and P2 from salt fractionated whole testes derived from WT, AA, and EE mice. **(C)** Violin plots of initial (left) and residual (right) compaction after IDC1 and IDC3 for each protamine construct. **(D)** Example traces showing P1-DNA decompaction steps for AA and EE from IDC1 and IDC3, data in grey and fitted steps in magenta and gold for AA and EE, respectively. **(E)** Joint histogram of AA-DNA decompaction steps from IDC1 and IDC3, grey dashed lines indicate peaks from Gaussian fits. **(F)** Joint histogram of EE-DNA decompaction steps from IDC1 and IDC3, grey dashed lines indicate peaks from Gaussian fits. **(G)** Kinetic parameters of step dwell times. (top) Pie charts showing populations of fast steps (lighter shade) and slower steps (darker shade). (bottom) Average step dwell rates for each protamine construct from IDC1 and IDC3.

To examine how these mutations affect protein stability *in vivo*, we repeated salt fractionation experiments on testicular spermatids from both WT and mutant mice. We observed that AA has similar salt fractionation dynamics to WT P1, including a substantial portion of AA protein that persists in the insoluble fraction (**Figure 5B**). In contrast, EE is markedly less stable and very little of the mutant protein remains in the insoluble fraction (**Figure 5B**). Interestingly, this instability also affected P2 retention in EE spermatids, where a portion of P2 dissociated at lower salt concentrations—a phenomenon not observed in salt fractions of WT or AA spermatids (**Figure 5B**). These findings indicate that the EE mutations also impair P2 protein stability and chromatin binding *in vivo*.

### The charge status of P1 S43/T45 tunes DNA binding affinity and compaction mechanics

Because of the altered stability of P1-DNA complexes caused by substitution of S43 and T45 to either alanine or glutamic acid, we returned to optical tweezers to test the effects of these mutations on P1-DNA structure formation and disassembly. These experiments were performed as described above for WT P1 (**Figure 1A**). During the incubation phase, we find all three proteins exhibit a force creep, working against the trap to compact DNA (**Figure 1B (WT), S5G (AA,EE)**). The force creep for WT P1 and AA both plateau at ≈7pN, while EE levels off at a slightly lower force of ≈6pN, in line with the EMSA and anisotropy results (**Figure 1B, S5G**). Furthermore, we measure the amount of DNA compacted by each mutant at the end of the incubation phase (initial compaction) and find that both WT P1 and AA package roughly the same amount of DNA during the incubation across IDCs, while EE induces ≈35% less initial compaction (**Figure 5C**). Moreover, by examining the residual amount of compacted DNA remaining at the end of the decompaction phase, we find that in IDC1—which occurs with protamine in the buffer—both WT P1 and AA have a sizable population of residual compacted structures (**Figure 5C**). However, almost all initially compacted structures formed by EE in the first IDC decompacted completely (**Figure 5C**). Across P1 conditions, the differences in compaction levels largely disappear by the third IDC (**Figure 5C**). Together, these data agree with the conclusion that glutamic acid substitutions destabilize P1-DNA structures and also indicate an impaired ability to form structures.

Throughout all IDCs we find that both AA and EE exhibit stepwise decompaction (**Figure 5D-F, S5H,I**). In IDC1 and IDC3, the step size distribution for AA exhibits peaks at 40bp, 80bp, and 120bp (**Figure 5E**). As observed for WT P1, the relative frequency of larger structures decreases from IDC1 to IDC3, and AA traces display compaction steps with considerable variance in magnitude (**Figure S5I**). However, in contrast to WT P1, the sizes of AA decompaction steps follow a trend of integer multiples (40xN bp) and sizes of AA-DNA structures are largely the same in both IDC1 and IDC3 (**Figure 5E**). Comparatively, the step size distribution for EE in IDC1 displays peaks at approximately 50bp and 95bp, which then reduce in IDC3 to 40bp and 80bp (**Figure 5F**). EE also exhibits compaction steps, but generally exhibits fewer steps across all conditions, likely owing to compromised DNA interactions (**Figure S5I,K**). Notably, ≈80bp steps appear in at least one IDC for all three P1 proteins, suggesting this size may be energetically favorable enough to overcome other factors dictating structure formation (**Figure 1D, 5E,F**).

Next, we examined the step dwell times of the P1 phosphorylation mutants. In IDC1, when protamine is present during compaction, AA decompaction steps are well described by a single rate (**Figure 5G, S5J,K**). However, in IDC3, a mixed population of relatively weaker and stronger states emerges (**Figure 5G, S5J,K**). This indicates that when AA is abundant, it creates more homogeneous P1-DNA structures, possibly due to competition between free and bound protamine that limits the strengthening or formation of more stable assemblies. Alternatively, EE displays at least two dissociation rates across all IDCs (**Figure 5G, S5J, K**). While the stronger state (i.e., the slower rate) in EE resembles the slower rates for WT P1 and AA, its faster rate is twice that of WT P1 and AA (**Figure 5G**). Nominally, across all P1 conditions, the relative number of weak steps is ≈3 times higher (**Figure 5G, S1G**,**S5K**), indicating that the overall decompaction behavior *in vitro* is largely determined by the weaker population. Overall, these results suggest that altering the charge status of S43 and T45 modulates the size and stability of P1– DNA structures, and that the relative abundance of free P1 can shift the balance between protamine species to favor or disfavor particular structural populations.

### Loss of P1 S43/T45 phosphorylation compromises developmental competency

We next asked whether the distinct effects of the AA and EE substitutions on P1-DNA compaction stability translate to reproductive fitness and embryo developmental competence. Fertility was first assessed by natural mating, a composite measure reflecting sperm motility and fertilization efficiency. In this assay AA males produced litters (albeit slightly smaller litter sizes; WT (≈9.5pups/litter); AA (≈7.5pups/litter), whereas EE males were sterile (no pups), consistent with their severe sperm motility defects (**Figure 6A**). To directly assess fertility independent of motility or fertilization barriers, we performed intracytoplasmic sperm injection (ICSI) using oocytes collected from the same pool of WT females. ICSI experiments revealed that although both mutants experienced a modest delay in progression to the 2-cell stage at 18 hours post fertilization (hpf), embryos derived from EE sperm had a marked developmental arrest at the 2-cell stage (**Figure 6B**). As a result, blastocyst formation was significantly compromised when compared to WT controls (4% vs. 63%). Notably, despite the relatively normal fertility observed for AA mutants in natural mating, fewer embryos reached the blastocyst stage (40% vs. 63%), with losses distributed across all stages of pre-implantation development.

**Figure 6:**
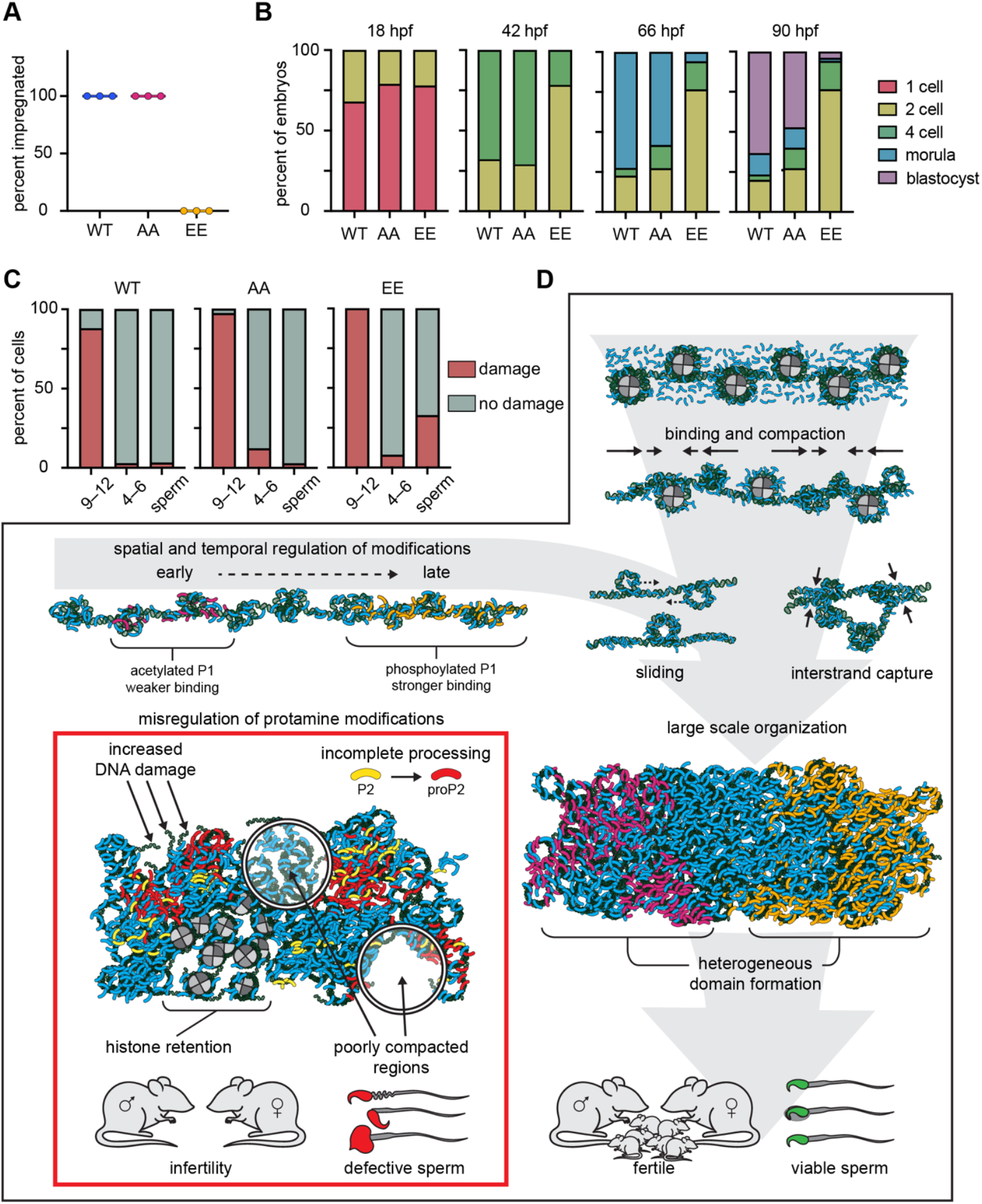
Loss of P1 S43/T45 phosphorylation compromises developmental competency. **(A)** Fertility outcomes from WT, AA, and EE males quantified as percent of females impregnated. Each dot represents a single male, and each male was mated to n=3 independent females. **(B)** ICSI developmental outcomes by day as the percent of embryos in each stage. For hours post fertilization (hpf) 42 and beyond, 1-cell stalled embryos were excluded for simplicity. Data were collected from n=2 males per genotype. **(C)** DNA damage levels present in stage 9–12 spermatids, stage 4–6 spermatids, and mature sperm across genotypes determined by a neutral Comet assay. Data were collected from n=2 males per genotype. **(D)** Model illustrating the spatial and temporal organization of P1 modifications and their impact on sperm chromatin architecture and fertility. During normal spermiogenesis, P1 molecules acquire distinct PTMs in a temporally regulated manner, generating populations with variable charge and DNA-binding strength. P1 binding induces chromatin compaction and drives the formation of multiple local DNA loops that subsequently coalesce through sliding and inter-strand capture to create large-scale, compact, and mechanically stable chromatin domains. Protamine PTMs modulate each stage of this condensation program. The differential localization and abundance of individual P1 PTMs establish heterogeneous yet ordered compartments within mature sperm, a pattern that is critical for proper chromatin organization and subsequent embryonic development. However, the loss of modifications (inset) alters the ratio of modified P1 species and the P1:P2 balance, leading to incomplete protamine exchange, deficient processing of P2, retention of histones, DNA damage, and poorly compacted chromatin. These structural defects result in abnormal sperm morphology, genomic instability, and impaired fertility.

Given the pronounced block at the 2-cell stage for EE-derived embryos, we next asked whether elevated DNA damage might underlie this phenotype. DNA damage is known to compromise embryo development by delaying DNA replication in the zygote and impairing 2-cell progression.^48^ During normal spermiogenesis, programmed DNA double-strand breaks (DSBs) are introduced in early elongating spermatids but typically resolve as chromatin compaction proceeds. Thus, DNA damage detected in mature sperm could arise either from unresolved DSBs formed during spermiogenesis or from new breaks accumulated during epididymal transit as a result of insufficient chromatin condensation. To distinguish between these possibilities, we monitored the dynamics of DNA damage in spermatids isolated from different stages of seminiferous tubules using either a Comet assay (**Figure 6C**) or gH2AX staining **(Figure S6A,B**): IX-XII (actively undergoing programmed DSBs), IV-VI (which should have completed repair), and in epididymal sperm. As expected, DNA damage is highest in WT spermatids isolated from stages IX-XII (**Figure 6C, S6A,B**), but most spermatids have repaired DSBs by stages IV-VI. Similar to WT, spermatids from stages IX-XII in both mutants had the highest fraction of DSB+ cells, but unlike WT, a higher fraction of cells had persistent DNA damage in stages IV-VI. These lingering breaks were largely resolved in AA males by the mature sperm stage. In contrast, EE spermatids appear to acquire additional damage during spermatid maturation leading to roughly ≈30% of mature sperm exhibiting damage as compared to ≈3-4% of WT and AA sperm (**Figure 6C**). We then turned to a TUNEL assay which, unlike the Comet assay, allows us to correlate DNA damage with nuclear morphology. Given the heterogeneity in sperm head sizes observed in EE males (**Figure 4F**), we quantified TUNEL signal intensity as a function of nuclear area. This analysis revealed that larger, more decondensed sperm nuclei exhibited higher levels of DNA damage, demonstrating that decompacted sperm cells are more prone to DNA damage (**Figure S6C,D**). Together, these results indicate that constitutive P1 phosphorylation compromises sperm chromatin integrity and reduces compaction, rendering the paternal genome more susceptible to damage during sperm maturation. These observations are consistent with our *in vitro* findings that the EE mutant binds DNA more weakly and forms less stable P1-DNA structures. This molecular instability likely underlies both the heterogeneous sperm morphology and the developmental arrest observed following fertilization.

## DISCUSSION

### P1-driven DNA condensation

As a consequence of protamine’s ability to compact DNA, it creates tension along the polymer. *In vitro*, we observe this phenomenon during the incubation phases of our experiments, where we show that WT P1 can induce up to ≈7pN of tension along a single DNA tether (**Figure 1B**). For context, the outer turn of the nucleosome unwraps when subjected to 3–6pN of external force, while the inner turn typically unwraps at forces greater than 15pN.^49,50^ Thus, protamine activity alone may be sufficient to liberate and package half of the DNA organized by nucleosomes, while also significantly lowering the barrier to release the remaining histone-bound nucleotides. The implication is that protamine does not passively rely on cellular machinery but actively participates in the global histone-to-protamine exchange. In addition to linear compaction, protamine is also capable of capturing and compacting DNA in three dimensions.^51-53^ In this work, we demonstrate that an extended DNA fiber decorated with protamine can capture additional dsDNA molecules from their local environment (**Figure 2B**). Moreover, we show that captured DNA serves as a scaffold to attract more protamine, which leads to increases in mechanical strength and stability (**Figure 2F**). Unexpectedly, during the initial period after association, captured DNA exhibits diffusive behavior (**Figure 2E**). As captured DNA slides along protamine-bound DNA, it can combine with existing or nucleate new P1-compacted structures (**Figure 2C,D**). Altogether these results support a model where DNA-associated proteins are subjected to pervasive destabilizing forces while the mechanical stability of chromatin continuously increases as protamine progressively packages the genome during sperm formation (**Figure 6D**). Importantly, this model does not necessitate that organization proceeds linearly along the genome, but, instead, may progress locally, engaging multiple distal regions of the same or separate chromosomes. Moreover, the finding that P1-bound DNA molecules can slide along one another provides a mechanism to both align specific regions of the genome during packaging, and to loop out intervening stretches of DNA between P1-DNA structures to allow packaged elements to achieve ideal densities or to reorganize into energetically preferred structures.

### P1 PTMs are temporally controlled

The protamine PTMs, K49ac and S43/T45ph, are established in different stages across the seminiferous epithelial cycle, which argues for differences in function. We show that K49ac is established early in spermiogenesis and persists at low levels in mature sperm (∼10% of P1), whereas S43/T45ph is established in later stage spermatids and is retained in mature sperm at much higher levels (∼40% of P1) (**Figure 3B,C,E, S3C**). Furthermore, our data reveals that phosphorylation of P1 plays a critical role in optimizing sperm chromatin for development, however, the temporal regulation of this modification may possibly be more important than its actual presence. This is supported by the observations that substitution of S43 and T45 for unmodifiable alanine results in subtle defects in sperm chromatin packaging and embryonic development, whereas substitution for glutamic acid (to mimic premature and constitutive phosphorylation) results in total loss of fertility, abnormal retention of histones, incomplete processing of P2, sperm DNA damage, and severely compromised embryonic development (**Figure 4G,H, 6A,B,C**). Together, these findings point toward a temporally controlled regulatory role for protamine PTMs that ensures proper sperm chromatin formation and early embryonic development (**Figure 6D**).

### P1-DNA interactions are inherently heterogeneous in vitro

Overall, we find heterogeneity *in vitro* in both the size and strength of compacted DNA structures made by protamine. By subjecting protamine-compacted DNA to elevated forces, we observed decompaction in short, discrete steps (**Figure 1C**). The presence of stepwise decompaction suggests that P1 forms small-scale, distinct structures when organizing DNA. An obvious candidate for a structure that agrees with our observations is a loop, or series of combined loops, and, notably, protamine has been directly shown to form loop structures when compacting DNA *in vitro*.^54-57^ Here, we found that the length of DNA released when P1-DNA structures relax under force ranges from 30 to 150bp, and larger structures tend to be more mechanically stable than smaller ones (**Figure 1D, S1H**). Together, this suggests that compacted DNA structures are likely organized by multiple P1 molecules that individually contribute to overall strength, and the release of compacted DNA is cooperative under tension. Interestingly, for both WT P1 and EE, decompaction steps are not integer multiples of a fundamental size (**Figure 1D, 5F**). This heterogeneity in sizes may be an inherent property of how protamine compacts DNA, or due to the diversity of PTMs within our natively derived P1 samples. Notably, phosphorylated protamine is expected to have a slightly smaller footprint on DNA than unmodified protamine.^58^ As a result, incorporation of phosphorylated P1 within a structure may alter the size relative to structures containing the same number of unmodified P1 molecules. Consistent with this interpretation, we show that the non-phosphorylatable AA protein exhibits regular decompaction steps that follow a clear size trend, suggesting a more homogeneous makeup of compacted structures (**Figure 5E**).

When examining the decompaction rate of individual P1-DNA assemblies, we found evidence for two structures: a slowly decompacting, stronger state and a faster relaxing state more susceptible to disruption. While all three of the P1 proteins tested here formed both stronger and weaker complexes with DNA, the stronger state was largely unpopulated in conditions where AA was in excess (**Figure 5G**). This suggests that competition between AA molecules for binding sites inhibits either the formation of stronger structures or the transition from weaker to stronger states. However, the inclusion of phosphorylated or negatively charged C-terminal residues appears to override this competition. This is especially surprising because EE generally binds weaker to DNA, as evidenced by binding anisotropy, force creep, and decompaction data, yet even under competitive conditions, it is still able to create stronger structures (**Figure 1B, 5A,G**). Together this points to a role for P1 phosphorylation in stabilizing compacted DNA but also argues that phosphorylation timing and levels should be well controlled as not to overly inhibit initial binding and compaction. Despite a clear difference in the *in vitro* binding properties of P1 proteins tested here, it remains surprising that P1-DNA interactions, which are presumably driven by arginine-DNA electrostatics, should be sensitive to the small changes that PTMs induce. However, our results argue that these changes must occur in locations related to the specific structures made between P1 and DNA or between protamine molecules.

### P1 PTMs create distinct chromatin states in vivo

We also observe evidence for protamine-based heterogeneity *in vivo*. Both P1 and P2 exhibit different chromatin association strengths and phosphorylated, but not acetylated P1, occupies overlapping domains with P2 in sperm (**Figure 1F,G S3G**). We show that these different states are driven not by P1 and P2 themselves but are instead epigenetically governed by PTMs on P1. Importantly, these modifications drive distinct nuclear localization patterns. Specifically, K49ac exhibits a punctate staining pattern in both spermatids and mature sperm, suggesting that this modification may be present programmatically in specific regions or loci (**Figure 3F**). P1 S43/T45ph, on the other hand, is more diffuse yet shows clear enrichment for the nuclear periphery, consistent with previous work showing that phosphorylated P1 interacts with Lamin B receptor at the nuclear periphery (**Figure 3F,S3G**).^59^ Furthermore, these modifications exhibit significantly different association strengths to chromatin. K49ac remains weakly associated both in spermatids of the testis and mature sperm, whereas S43/T45ph is strongly associated with chromatin in both cell types (**Figure 3G**). How these different chromatin states relate to the weak and strong states we observe *in vitro* will require further investigation. However, the clear partitioning of these modifications we observe both histologically and biochemically supports our revised model of sperm chromatin wherein P1 modifications establish distinct chromatin states (**Figure 6D**).

### The role of heterogeneity

Heterogeneity within the protamine-bound genome may serve several functions. Broadly speaking, heterogeneous interactions between protamine and DNA driven by PTMs provide a mechanism to affect differential packaging of the sperm genome. Because retained histones make up only 1-5% of mouse sperm chromatin, heterogeneity within protamine-bound regions (accounting for 95-99% of the genome) provides a greater canvas for a large-scale control of unpackaging after fertilization. Salt fractionation of both testicular spermatids and mature sperm revealed that regions harboring K49ac are not as stably compacted as regions containing S43/T45ph. Therefore, it is possible that these modifications, because of their biochemical properties on chromatin, instruct the programmed timing of genome unpackaging in the zygote. Alternatively, specific genomic regions may recruit factors based on the proportions of P1 PTMs and/or biophysical differences (stronger vs. weaker states) of PTM-driven P1-DNA compaction. Such a system is also consistent with the rapid evolution of held notion that protamine-based chromatin is inherently homogeneous, and that sperm contributes no information to the zygote beyond the information held within retained histones and retained DNA methylation. Instead, we propose a model wherein protamine PTMs serve as species-specific epigenetic carriers and create heterogeneous states within the protamine-bound genome (**Figure 6D**). Ultimately, this code fine-tunes sperm reproductive fitness by ensuring mechanical protection prior to fertilization and controlled unpackaging in the zygote.

## ACKNOWLEDGEMENTS

We thank members of the Hammoud and Redding labs for scientific discussion and manuscript feedback. We acknowledge Thomas L. Saunders, Zachary T. Freeman, Wanda Filipiak, Galina Gavrilina, Honglai Zhang, Eden Dulka, Jennifer Leo, and Kay Oravecz-Wilson of protamines across species and the lineage specificity of P1 C-terminal modifications. In other words, not enough evolutionary time exists to create an entirely new reader/writer system for emergent modifications; however, our model supports the notion that the sequence grammar of the protein primes its epigenetic properties to fine-tune chromatin structure in a species-specific manner.

*In vivo*, likely both the biophysical changes brought on by PTMs and otherwise direct interactions between protamine PTMs and other proteins gate proper loading, unloading, and remodeling of the protamine organized genome. It is possible that loading of protamines is influenced by the already present epigenetic state of chromatin, and protamine PTMs are preferentially loaded in ways that preserve this information in mechanical detail, implying that the somatic epigenetic code directs the heterogeneous landscape of developing sperm. Additionally, protamine heterogeneity may also offer genomic access in poorly or weakly compacted regions for the recruitment of other factors.^60^ Alternatively, there may be yet downstream remodeling events that re-organize disordered protamine-chromatin interactions to create a more uniform packaged ultrastructure. Altogether, our work challenges the previously the Transgenic Animal Model Core of the University of Michigan’s Biomedical Research Core Facilities for design and production of P1-V5 transgenic mice; Yu-Chiang Hu and members of the Cincinnati Children’s Hospital Transgenic Animal and Genome Editing Core Facility for generation of P1^S43AT45A^ and P1^S43ET45E^ mice. Research reported in this publication was supported by National Institutes of Health (NIH) grants 1DP2HD091949-01 (S.S.H.), R01HD104680 01 (S.S.H. and S.R.), R01HD113274 (S.S.H.), GM148028 (S.S.H), R35GM141736 (S.R.), 5R03HD101501 (S.B.S), 1F31HD117648-01 (C.A.T.), training grants T32-HD079342 (S.S.), T32GM007544 (C.A.T.), and Open Philanthropy Grant 2019-199327 (5384) (S.S.H.).

## AUTHOR CONTRIBUTIONS

L.M., C.J.W., S.R., and S.S.H conceived and designed the experiments. Experiments were 20 performed by L.M., C.J.W, R.A., M.R., S.B.S., W.X., C.A.T., S.S., and Y.S. M.R.B analyzed fluorescence anisotropy data. Technical oversight was provided by P.J.O., R.A.H., and K.E.O. The manuscript was written by L.M.,

## METHODS

### Lead contact and materials availability

Requests for resources and reagents and additional information should be directed to and will be fulfilled by the lead contact, S.S.H. All unique reagents generated in this study are available with a completed Materials Transfer Agreement. All optical tweezer analysis, including python notebooks containing all code used to create all figures presented in this manuscript, and experimental data is accessible at: https://uma1.osn.mghpcc.org/sy-redding-bmb/lm-cw-2025-data (DOI: 10.5281/zenodo.17554687).

### Mice

All experiments using animals (*Mus musculus*) were carried out with prior approval of the University of Michigan Institutional Animal Care and Use Committees of the University of Michigan (Protocols: PRO00006047, PRO00008135, PRO00010000, PRO00011691) and in accordance with guidelines established by the National Research Council Guide for the Care and Use of Laboratory Animals. Mice were housed in the University of Michigan animal facility in an environment controlled for light (12 hours on/off), temperature (21-23°C), and humidity (30-70%) with ad libitum access to food and water (Lab Diet no. 5008 for breeding mice, no. 5LOD for non-breeding mice). P1^S43AT45A^ and P1^S43ET45E^ mice were generated on the C57BL/6N background using CRISPR-Cas9 mediated genome editing by the Cincinnati Children’s Hospital Transgenic Animal and Genome Editing Core Facility as previously described.^43,61^ Animal procedures were carried out with prior approval of Cincinnati Children’s Hospital Medical Center and in accordance with the Institutional Animal Care and Use Committee. All mice were backcrossed to the C57BL/6J background. All mice used for experiments were males between 8-20 weeks of C.J.W., S.R., and S.S.H. with input from all authors.

## DECLARATION OF INTERESTS

The authors declare no competing interests. age. Adult female mice were used solely for breeding purposes.

### Antibodies

A rabbit polyclonal antibody against P1 S43/T45ph was generated by LabCorp Drug Development (formerly Covance) via immunization of rabbits with the following peptide: CRRRRS(p)YT(p)IRCKKY. The P1 S43/T45ph antibody was used at 1:1000 for immunoblotting and 1:500 for immunofluorescence. All other antibodies used in this study are specified below and were used at the indicated dilutions: P1 (Briarpatch Biosciences, Hup1N): 1:500, P2 (Briarpatch Biosciences, Hup2B): 1:1000, P1 K49ac^43^: 1:500, PNA-Lectin (Genetex, GTX01508): 1:1000, histone H2B (Abcam, ab52484): 1:1000, histone H3 (Abcam, ab1791): 1:1000, TNP1 (ProteinTech, 17178-1-AP): 1:100 for immunofluorescence, 1:500 for immunoblotting, TNP2 (Santa Cruz, 393843): 1:1000, V5 (BioRad, MCA1360): 1:1000, acH4 (Millipore, 06-866):1:1000, alpha tubulin (66031-1-Ig): 1:1000.

### Preparation of DNA reagents

For use in IDC experiments with free DNA, pUC19 plasmid was digested with HindIII restriction endonuclease at 37°C overnight. The reaction was then heat inactivated at 80°C for 20 minutes. Digested DNA was ethanol precipitated, resuspended in 1xTE buffer (10mM Tris-HCl pH = 8.0, 1mM EDTA), and quantified by absorbance at 260nm. For experiments using fluorescent pUC19, 35.14µg of digested pUC19 and 34.5U of T4 polymerase were incubated in 150µL of 1x NEB buffer r2.1 for 16 hours at 11°C without nucleotides to allow polymerase exonuclease activity to reveal small 5′ overhangs. Then 3µL of 6mM dATP, dCTP, and dGTP as well as 3.5µL of 1mM Alexa Fluor 647 dUTP were added. The reaction was then incubated for 30 minutes at 37°C, followed by 2 hours and 20 minutes at room temperature. Then 8.1µL of 2mM dTTP was added and the reaction was incubated for an additional 1 hour and 20 minutes at 11°C. The reaction was then heat inactivated at 75°C for 20 minutes. Finally, the DNA was purified using a Qiagen PCR Cleanup Kit (Cat no. ID 28106) and the concentration was determined by absorbance at 260nm and 655nm.

### Optical Trap Setup

The objective and condenser of the optical trap were aligned following manufacturer’s instructions. In all experiments, the temperature was set to 28°C, and Immersol™ W (Zeiss 444969-0000-000) was used on the objective instead of water. IR Lasers were set to the following for all experiments: 100% trapping laser power, 30% overall power, 50% split, and the brightfield LED was set to 50%. Prior to each experiment, the flow cell channel was washed with 0.5mL of H2O, then washed with 0.5mL of 1mg/mL BSA, and finally 0.5mL of 0.5% w/v Pluronic® F-127. Next, all syringes were washed with 0.5mL of experimental buffer (40mM Tris-HCl pH 7.65, 50mM NaCl, 5mM MgCl_2_, 5mM DTT, 1mg/mL BSA, passed through a 0.22µm filter) to clear any residual Pluronic® F-127. Then, each fluidic channel was loaded with the proper reagent for the experiment. Each channel contained 0.5mL of experimental buffer. In addition, 5µL Streptavidin-coated Ø4.0-4.9 µm beads (Spherotech, SVP-40-5) were added to channel 1, 10ng biotinylated lambda DNA (Lumicks, SKU:00001) was added to channel 2, and protamine 1 (P1) was added to channel 5 to 150nM. In experiments containing free pUC19 dsDNA, pUC19 was added to channel 4. Note, protamine stocks were incubated at 37°C for 15 minutes and spun down on a benchtop microcentrifuge for 1 minute prior to pipetting P1 into room temperature experimental buffer. Next, all flow channels were opened and allowed to flow at 0.4 bar of pressure for two minutes. Then, flow was terminated, and force calibration was performed via the thermal/passive calibration method. Finally, the piezo distance setting was turned on and a python script automating the entire experiment was executed.

### Experimental Workflow of Incubation Decompaction Cycle Experiments (FCv6 Script)

First, two beads tethering one molecule of biotinylated lambda DNA was acquired under flow. Next, we verified that only one DNA was tethered between the beads by performing a control pull in channel 4 (buffer only). Then, we perform an incubation decompaction cycle (IDC) in channel 4. In each IDC the tether is incubated at an extension of 6µm for 180 seconds, then an 8pN force clamp is applied for 150 seconds, finally the tether is returned to an extension of 6µm (**Figure 1A**). Data is recorded during the incubation, force clamp, and return. After the control IDC (in buffer), the DNA tether is moved into channel 5 containing P1, and IDC1 is performed. After IDCI, we then perform IDCII, but, while the incubation occurs in channel 5 (with P1), the tether is moved into channel 4 (buffer) before the 8pN force clamp is applied. Then, IDCIII is performed exclusively in channel 4 (buffer). Finally, the beads are discarded and the experiment restarts.

### Experimental Workflow of Kymogram Collection (FCv6dFluor Script)

The setup for fluorescent DNA capture experiments was identical to the FCv6 method except either 3.6pM or 18 pM of pUC19-HindIII AF 647-dUTP (AF647 DNA) was added to channel 4 (**Figure 2A**). These experiments begin, as above, by acquisition of a single DNA tether. Then, the brightfield LED was turned off and the 638nm laser was set to 2%. Next, the DNA tether was incubated in channel 4 (AF647 DNA) for 60 seconds at an extension of 6µm before extending to 15.25µm. Then a time series of 10 images were collected (**Figure S2A**). Next, the tether was moved into channel 5 (P1), incubated at an extension of 6µm for 60 seconds and then extended to 15.25µm, and again imaged for 10 frames. Then, the tether is moved back into channel 4 (AF647 DNA), incubated and extended as previously, but then imaged in kymogram mode for 10 minutes (**Figure 2C,D**). Finally, the beads are discarded, and the experiment restarts.

### Experimental Workflow of Incubation Decompaction Cycle Experiments with Free DNA (FCv7 Script)

The setup for unlabeled DNA capture experiments was identical to the FCv6 method except for conditions involving incubation with free DNA where 0.21nM pUC19-HindIII (dsDNA) was added to channel 4. As above, a single DNA tether was captured and verified. Next a control incubation decompaction cycle was performed in channel 4 (dsDNA or buffer). In each IDC the tether was incubated at an extension of 6µm for 180 seconds, then an 8pN force clamp was applied for 150 seconds, and finally the tether was returned to an extension of 6µm. Data were recorded during the incubation, force clamp, and return. After the control IDC, the tether was moved into channel 5 (P1), and an IDC was performed. We then perform slightly altered IDC. First, the tether was incubated in channel 5 (P1) at an extension of 6µm for 180 seconds, then the tether was moved to channel 4 (dsDNA or buffer) and incubated at an extension of 6µm for 180 seconds before then moving back into channel 5 (P1) to incubate again at an extension of 6µm for 180 seconds. Following these three incubations, we then apply an 8pN force clamp for 150 seconds (**Figure S2B**). Then the tether is returned to channel 4 (dsDNA or buffer) and incubated again at 6µm for 180 seconds before applying an 8pN force clamp for 150 seconds (**Figure 2F**). Finally, the beads are discarded, and the experiment restarts.

### Optical trap experiment analysis

For each pulling experiment we collect four trajectories: a DNA control pull followed by three IDCs (**Figure 1A**). First, the control pull of each tether is analyzed to ensure that it behaves as a single dsDNA molecule and does not exhibit residual tension at 6μm extension^62^. Any tether that does not meet these criteria are not included in downstream analysis.

#### Force Creep Analysis

For each DNA tether, there are four collected incubation phases: control and three IDC incubations. The force measurements during these incubation phases were down sampled to ≈78Hz and grouped by experimental phase. Then, grouped data were averaged and plotted as mean +/-standard error of the mean versus time and the data were fit to the complementary distribution function of an exponential decay distribution multiplied by a constant (**Figure 1B, S5G**).

#### Step finding

For each dataset, we first determine the mean tether length of the DNA in the control pull at the applied clamping force to assign an end position for the trace and calculate the per base pair extension for each DNA tether. To do this, we define the region of the data where the force clamp is stable (i.e., after the initial ramp up) by down sampling the measured force to 50Hz and applying a Savitzky Golay filter to both smooth the data and determine its first derivative. The derivative of the force becomes exceedingly small as the force clamp becomes stable (**Figure S1A,B**). We then take the average extension of the DNA molecule over this stable force regime as the mean tether length of the DNA. Using this value, we then filter out any traces with abnormal extension and determine the per base pair extension as the length in base pairs of lambda phage DNA (48502bp) divided by the mean tether length. Next, we analyze each IDC and determine stepwise events. In each case, we again truncate each trace to the region where the trap force is stable in the same manner as for the DNA control pull. We also truncate traces if they extend to within 300nm of the mean tether length as determined in the DNA control pull. Then, we resample the data to 15kHz, and determine the likely decompaction and compaction steps by applying a slightly modified version of the AutoStepfinder algorithm^63^ (**Figure 1C, 5D**). The output steps are then filtered by removing steps where the Z-score of the step assignment error is larger than one, this eliminates outliers only from the right-hand side (higher error) of the assignment error distribution. We also recover the step dwell times from the AutoStepfinder output and filter them as above (**Figure 1E, S5J**). In addition, for each IDC, we determine the mean initial and residual compaction levels as the average over the initial or final 100 position measurements from each truncated trace (**Figure 5C**).

#### Step size histograms

To ascertain the mean step sizes from the distribution of determined steps for each condition (i.e., IDCs, mutants), we fit each dataset to a Gaussian mixture model using publicly available libraries from Scikit-learn (Scikit-learn: Machine Learning in Python<http://jmlr.csail.mit.edu/papers/v12/pedregosa11a.html>‘_, Pedregosa *et al.*, JMLR 12, pp. 2825-2830, 2011.). For each dataset, we first determine the most likely number of Gaussians to fit (between one and ten) as the minimum of the Bayesian information criterion for each possible number. Then, once fit, we exclude any Gaussians from the joint distribution that represent less than 10% of the total population. This results in ≈10-15% of steps not represented in the joint distributions presented in figures 1D, S1C, 5E,F, S5H. However, all steps are included along with individual Gaussian fits in the complete histograms presented in figures S1E and S5I.

#### Step dwell analysis

Measured step dwells were separated into compaction and decompaction dwells by the sign of the associated step and we largely compute the rate parameters for only the decompaction steps. However, WT P1 compaction step data for each IDC is presented in Figure S1I. For each dataset, we use the bootstrap method to determine rate parameters and create the average plots presented in the figures. In all cases, we create 300 bootstrapped step dwell distributions, from which we then compute complementary cumulative distributions (survival functions). Then we take the average and standard deviation of these samples, which are plotted in the main and supplementary text in figures 1E, S1D,H,I, and S5J. Then we fit both a single and bi-exponential decay function to the first 99% of the data and compute the residuals to determine which model to use to best describe the data (**Figure S1F**). Once we determine the model, we then return to the bootstrapped sample distributions and fit each distribution individually with the appropriate model. The average and standard deviation of the fit parameters are then presented in figures 5G, S1G, and S5K.

#### Kymogram tracking and diffusion coefficient determination

The trajectories of bound fluorescent dsDNA were determined from kymograms by employing a tracking algorithm publicly available from the Pylake software library (https://lumicks-pylake.readthedocs.io/en/v1.6.2/tutorial/kymotracking.html). Diffusion coefficients were then determined from tracked DNA trajectories by performing a linear regression to the mean squared displacement (MSD) of each trace. We employed a previously described method to determine the ideal region of the MSD to perform the fit that minimizes error from correlated noise that arises in MSD curves.^46^ However, to keep the calculation of errors in longer traces computationally tractable, we down sample longer trajectories by interpolation to only include a maximum of 1000 data points. Importantly, due to the fractal nature of Brownian motion, this sampling has no effect on the determined diffusion coefficients.

#### DNA capture analysis

For experiments involving unlabeled dsDNA incubations, extension measurements are grouped by experimental phase: control or incubated with dsDNA, P1, or both. In each case, extension data were downsampled to ≈7.8Hz and normalized by setting their initial extension to zero. The extension values were then averaged and plotted as average extension +/-standard error of the mean vs. time. Finally, the data were fit to the complementary distribution function of an exponential decay distribution multiplied by a constant.

### Subcellular and high salt fractionation of protamines from adult testes and sperm

Germ cells from adult testes were subjected to subcellular and high salt fractionation as previously described.^64^ Briefly, frozen adult testes were homogenized using a Dounce homogenizer and spermatid tails were removed by treatment with 0.1% CTAB for 5 minutes on ice. Cells were then lysed for 15 minutes, centrifuged at 800xg for 15 minutes, and supernatants were removed as the cytoplasmic fraction. Nuclei were then first digested with 1 U of MNase at 37°C for 30 minutes before quenching with EGTA (supernatants kept as MNase fraction). Resulting chromatin was then fractionated via sequential treatments of 100 mM, 300 mM, 500 mM, 1 M, and 2 M NaCl for 30 minutes each. Proteins from all fractions were precipitated with 20% TCA overnight at −20°C before being used for immunoblotting.

For sperm, ∼30 million sperm were initially resuspended in 1 ml of 25 mM Hepes pH 7.5, 150 mM NaCl, 0.1% NP-40 to initiate cytoplasmic lysis before sperm nuclei were lysed in 50 mM Hepes pH 7.5, 150 mM NaCl, 0.05% L-alpha phosphatidyl choline, 0.5% Triton for 10 min on ice. Chromatin was then subjected to digestion with 5 U of MNase at 37°C for 30 minutes before quenching with EGTA (supernatants kept as MNase fraction). Resulting chromatin was then fractionated via sequential treatments of 100 mM, 300 mM, 500 mM, 1 M, and 2 M NaCl for 30 minutes each. Proteins from all fractions were precipitated with 20% TCA overnight at −20°C before being used for immunoblotting.

### Immunoprecipitations from adult testes

Adult P2^V5/+^ testes were homogenized in cell lysis buffer (15 mM Hepes pH 7.5, 15 mM NaCl, 25 mM KCl, 1 mM EDTA, 0.5 mM EGTA, 250 mM sucrose) using a Dounce homogenizer. Cells were then lysed by addition of NP-40 to a final concentration of 0.2% and incubated on ice for 15 minutes. Lysates were spun at 800xg for 15 minutes and the supernatant was set aside as the cytoplasmic fraction. Nuclei were resuspended in nuclear lysis buffer (50 mM Hepes pH 7.5, 2 mM EDTA, 2 mM MgCl_2_, 300 mM NaCl, 10% glycerol, 0.1% NP-40), sheared several times with a 23G needle, and incubated at 4°C for 4 hours with 1U of benzonase. Nuclear lysates were spun down at 10,000xg for 20 minutes and the resulting supernatant was combined with the cytoplasmic supernatant before adding to V5 beads. Lysates were incubated with V5 beads overnight at 4°C. Beads were then washed with 50 mM Hepes pH 7.5, 2 mM EDTA, 2 mM MgCl_2_, 150 mM NaCl and analyzed by immunoblotting.

### Acid extraction of sperm basic proteins

Acid extraction of protamines was performed as previously described.^43^ Briefly, sperm collected from the cauda epididymis and vas deferens were pelleted and subjected to hypotonic lysis in 1 mM PMSF. Sperm were spun at 8,000xg for 8 min and sperm pellets were resuspended in 100 mM Tris pH 8.0, 20 mM EDTA, 1 mM PMSF and then denatured for 10 min at room temperature, rotating, with 6 M guanidine-HCl, 575 mM DTT followed by alkylation with 522 mM sodium iodoacetate for 30 min in the dark. Protein pellets were washed twice with cold ethanol followed by extraction of basic proteins with 0.5 M HCl, 50 mM DTT at 37°C for 15 minutes. Supernatants were precipitated with TCA (20% final concentration) overnight at −20°C. The next day, precipitated proteins were spun down at 12,000xg for 10 minutes and protein pellets were washed twice with 100% acetone containing 1% BME before being resuspended in water.

### Peptide competition assay

To assess antibody specificity, protamines were first acid extracted from mature sperm as described above. For immunoblotting, antibodies were first incubated at room temperature for 30 minutes either alone or with tenfold excess of a specific or non-specific peptide. Blots were then incubated for 1.5 hours with antibody only or antibody with specific or non-specific peptide. The non-modified P1 peptide used was N-CRRRRSYTIRSKKY-C and the non-specific peptide used was N-DSNKEFGTSNESTE-C.

### Synchronization of spermatogenesis in juvenile male mice

Males were fed WIN 18,446, a retinoic acid inhibitor, daily beginning at 2 days post-partum (dpp) at a concentration of 100 μg/g body weight for 7 days.^65,66^ At 9 dpp, males were injected with 100 μg of retinoic acid (in DMSO) to initiate spermatogenesis. Testes were collected at 33 and 37 days after retinoic acid injection. Successful synchronization of spermatogenesis was confirmed by immunofluorescence of testes cross sections using DAPI and PNA-Lectin.

### Phenotypic assessment of P1^+/+^, P1^S43AT45A^, and P1^S43ET45E^ males

All phenotyping was performed on males at 9-12 weeks of age. Sperm were counted from n=3 technical replicates per animal. For fertility assessment, 8 week old males (n=3 per genotype) were individually housed for 3 days prior to addition of 8-week-old C57BL/6J females. Females (n=3 per male) were checked daily for the presence of copulatory plugs and placed in a new cage once a plug was observed to ensure only a single mating event.

### Neutral Comet assay for DNA damage

Comet assay to assess DNA damage in sperm and spermatids was performed using a neutral Comet kit (Trevigen) as previously described with modifications.^22^ 1-2 million sperm or ∼10,000 spermatids were collected from the epididymis, washed once with PBS, then mixed 1:10 with low melt agarose and spread onto slides. Sperm/agarose mixtures were allowed to solidify at 4°C for 30 minutes before being submerged in lysis solution for 1 hour. After 1-hour, fresh lysis solution containing 500 μg/ml Proteinase K was added and slides were incubated for 18 hours at 4°C. Lysis buffer was removed, and slides were washed with 1X neutral electrophoresis buffer for 30 minutes at room temperature. Slides were then subjected to electrophoresis in 1X neutral electrophoresis buffer at 1 volt per cm at 4°C for 45 minutes. Slides were then immersed in DNA precipitation solution for 30 minutes at room temperature, followed by washing in 70% ethanol (30 minutes at room temperature), dried, and stained with SYBR gold.

### Purification of protamines from mouse sperm and electrophoretic mobility shift assays

Protamines were purified from mature mouse sperm as previously described.^43^ Briefly, basic proteins were first extracted from mature sperm as described above, resuspended in 50 μl of water then brought up to 500 μl with 25 mM Hepes pH 7.5, 150 mM NaCl, 5 mM TCEP. Proteins were separated by size exclusion chromatography using a Superdex S75 column. Peak fractions were confirmed for the presence of P1 or P2 by immunoblotting and pooled prior to biochemical assays. For electrophoretic mobility shift assays, varying amounts of protamine were first incubated at 37°C for 10 minutes in reaction buffer before DNA (40 nM, 280 bp fragment) was added. Protamine-DNA complexes were then incubated at room temperature for 1 hour and resolved on a non-denaturing 0.5x TBE, 6% polyacrylamide gel. DNA was visualized using ethidium bromide and band intensities were quantified using ImageJ. For experiments in which P1 and P2 were mixed, P2 protein was purified from WT males and pro P2 was purified from EE males. P1 and P2 were mixed at a 1:2 ratio at the final concentrations depicted in Figure S5.

### Intracytoplasmic sperm injections

8-week-old females (B6D2F1/J) were superovulated using 7.5 IU of pregnant mare serum gonadotropin (Prospec Protein Specialists) ∼62 hours prior to oocyte collection, followed by injection of 7.5 IU of human chorionic gonadotropin (Sigma) ∼14 hours prior to oocyte collection. Gamete collection and piezo-actuated intracytoplasmic sperm injections were performed as previously described.^67^ Embryos were cultured in KSOM and monitored daily for developmental progress.

**Supplemental Figure 1:**
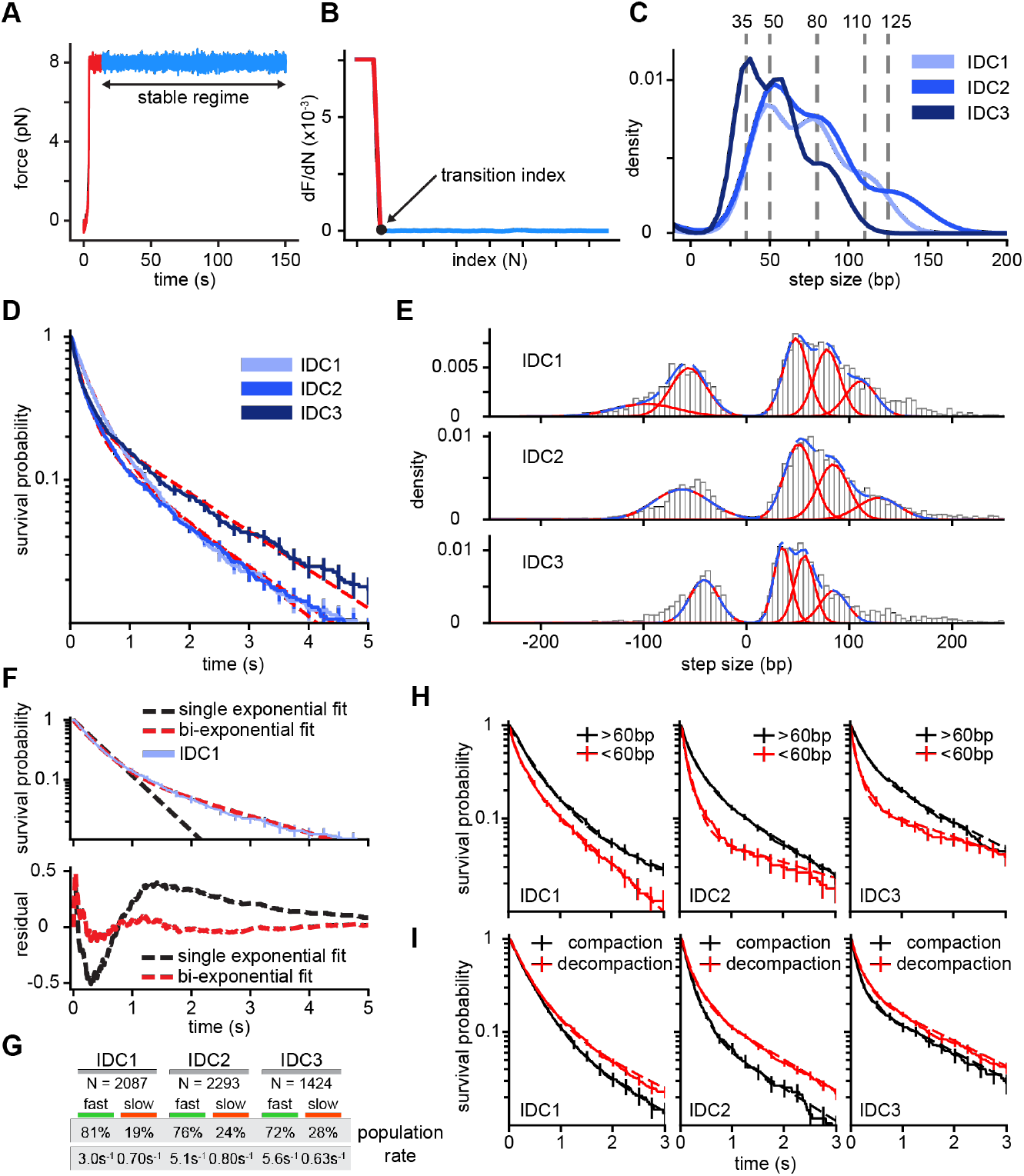
Protamine-DNA interactions organize into discrete structural units that undergo distinct decompaction kinetics. **(A)** Plot of force versus time for a sample DNA control pull. Full pull shown in red, stable force regime shown in blue. **(B)** Plot showing the first derivative of force. Black dot indicates the beginning of the stable regime. **(C)** Joint histogram of WT P1-DNA decompaction steps from all three IDCs, grey dashed lines indicate peaks from Gaussian fits. **(D)** Survival probability of step dwell times for WT P1-DNA decompaction steps from all IDCs, red dashed lines indicate bi-exponential fits. **(E)** Complete histograms of P1-DNA decompaction steps from each IDC. Individual Gaussian fits presented in red and combined fit in blue. **(F)** (Top) Survival probability of step dwell times for WT P1-DNA decompaction steps from IDC1. Dashed lines indicate single exponential (black) and bi-exponential (red) fits. (Bottom) Calculated residuals from each fit (colored as in top). **(G)** Kinetic parameters from fits to step dwell time survival probabilities for each IDC. **(H)** Survival probabilities of step dwell times for WT P1-DNA decompaction steps from all IDCs separated by size of associated step. Step dwells from steps less than 60bp in red, greater than 60bp in black. **(I)** Survival probabilities of step dwell times for WT P1-DNA from all IDCs separated by direction of associated step. Step dwells from decompaction steps in red, compaction steps in black.

**Supplemental Figure 2:**
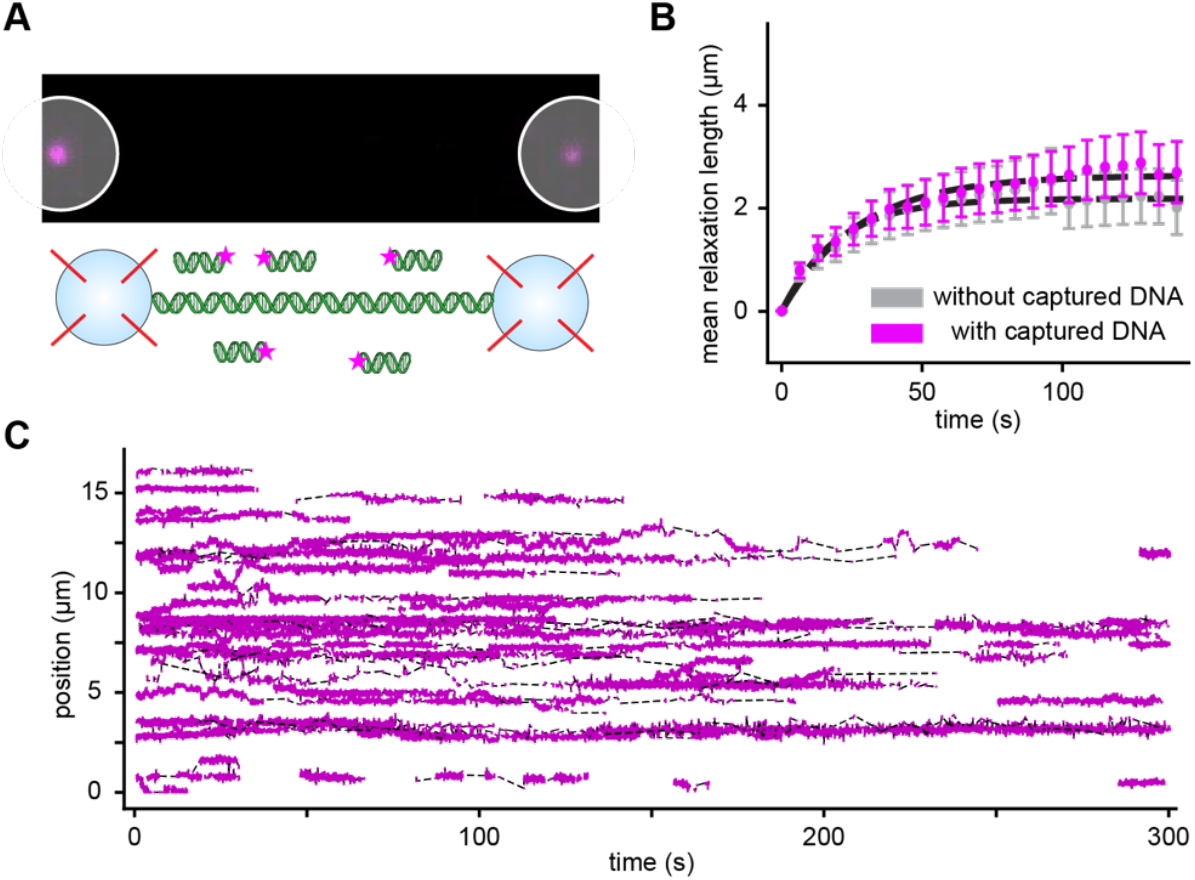
Interstrand captured DNA on P1-DNA complexes. **(A)** Example image (top) showing fluorescent DNA (magenta) is not captured when P1 is omitted from experiment. The underlying DNA tether is unlabeled, and beads are indicated by white circles, scale bar is 3μm. (Bottom) Cartoon of uncaptured DNA in solution. **(B)** Mean decompaction curves of DNA tethers in buffer containing WT P1. Tethers either previously incubated in buffer (grey) or 2.7kbp DNA (magenta), detailed experimental procedure described in methods. **(C)** Plot of all tracked captured DNA molecules used for diffusion coefficient determination.

**Supplemental Figure 3:**
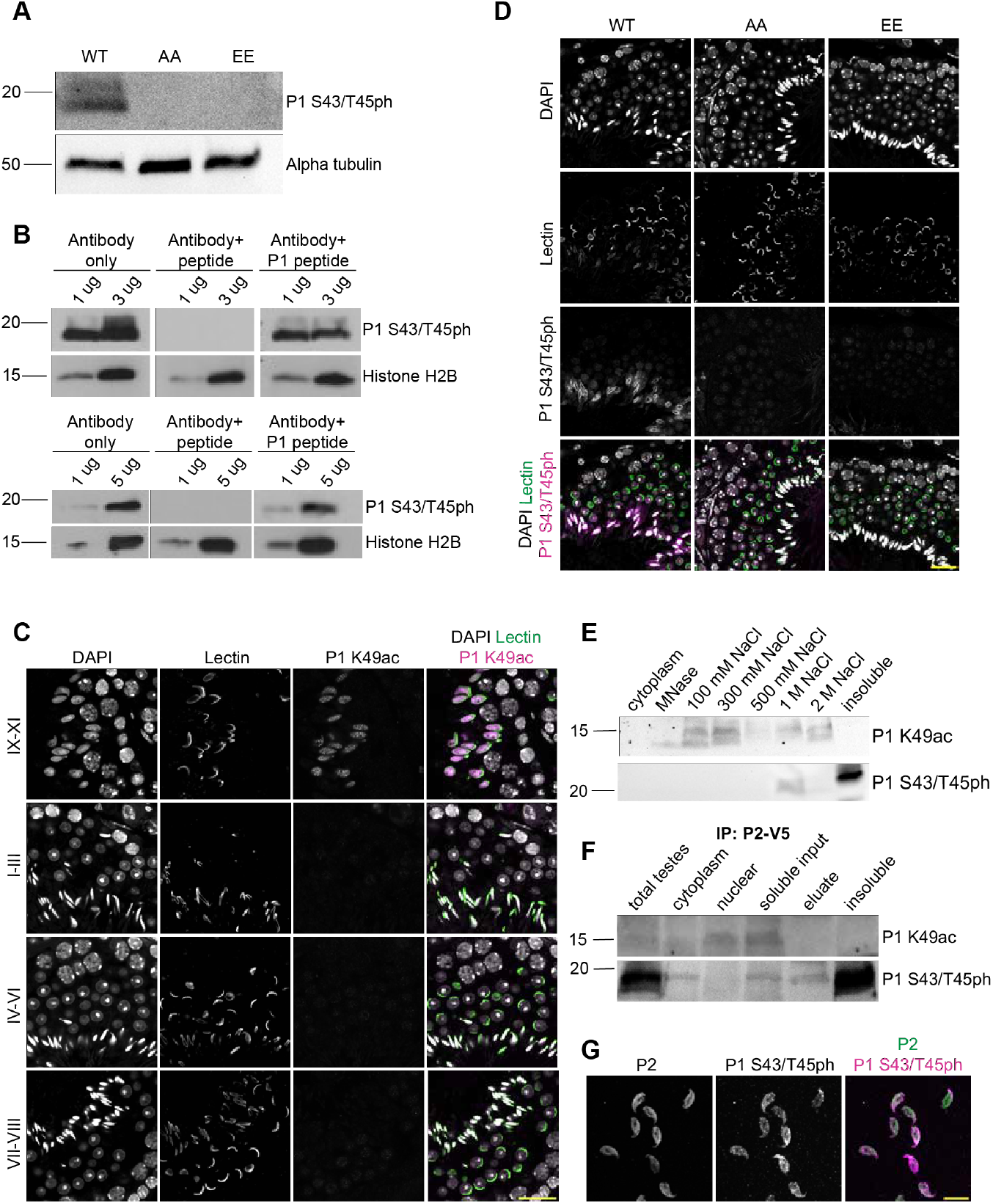
Sequential establishment of P1 K49ac and S43/T45ph generates protamine variants with distinct DNA-binding affinities *in vivo*. **(A)** Acid urea immunoblots of protein lysates from WT, AA, and EE tests illustrates specificity of the P1 S43/T45ph antibody as no band is detected from mutant lysates despite the presence of protein. **(B)** Immunoblot of sperm protein lysates shows a clear band for P1 S43/T45ph that is competed off only in the presence of a phospho-P1 peptide (top blot using an unmodified P1 peptide and bottom blot using a non-specific peptide). Shown are representative immunoblots, and similar results were obtained from n=2 independent experiments. **(C)** Immunofluorescence of adult testes cross sections stained for P1 K49ac across all stages of the seminiferous epithelial cycle highlights distinct patterns of stage specificity between K49ac and S43/T45ph. Scale bar: 20 μm. **(D)** Immunofluorescence of adult WT, AA and EE testes cross sections co-stained with the acrosome marker PNA-Lectin further confirms specificity of the P1 S43/T45ph antibody as no signal is detected in spermatids from either mutant. Scale bar: 20 μm. **(E)** Immunoblot of protein lysates from subcellular and high salt fractionated mature sperm probed for P1 K49ac or P1 S43/T45ph. Shown is a representative blot with similar results obtained from n=4 independent experiments. **(F)** Immunoblot of P2-V5 immunoprecipitated proteins probed for either P1 K49ac (top) or P1 S43/T45ph (bottom). Shown is a representative blot with similar results obtained from n=2 independent experiments.

**Supplemental Figure 4:**
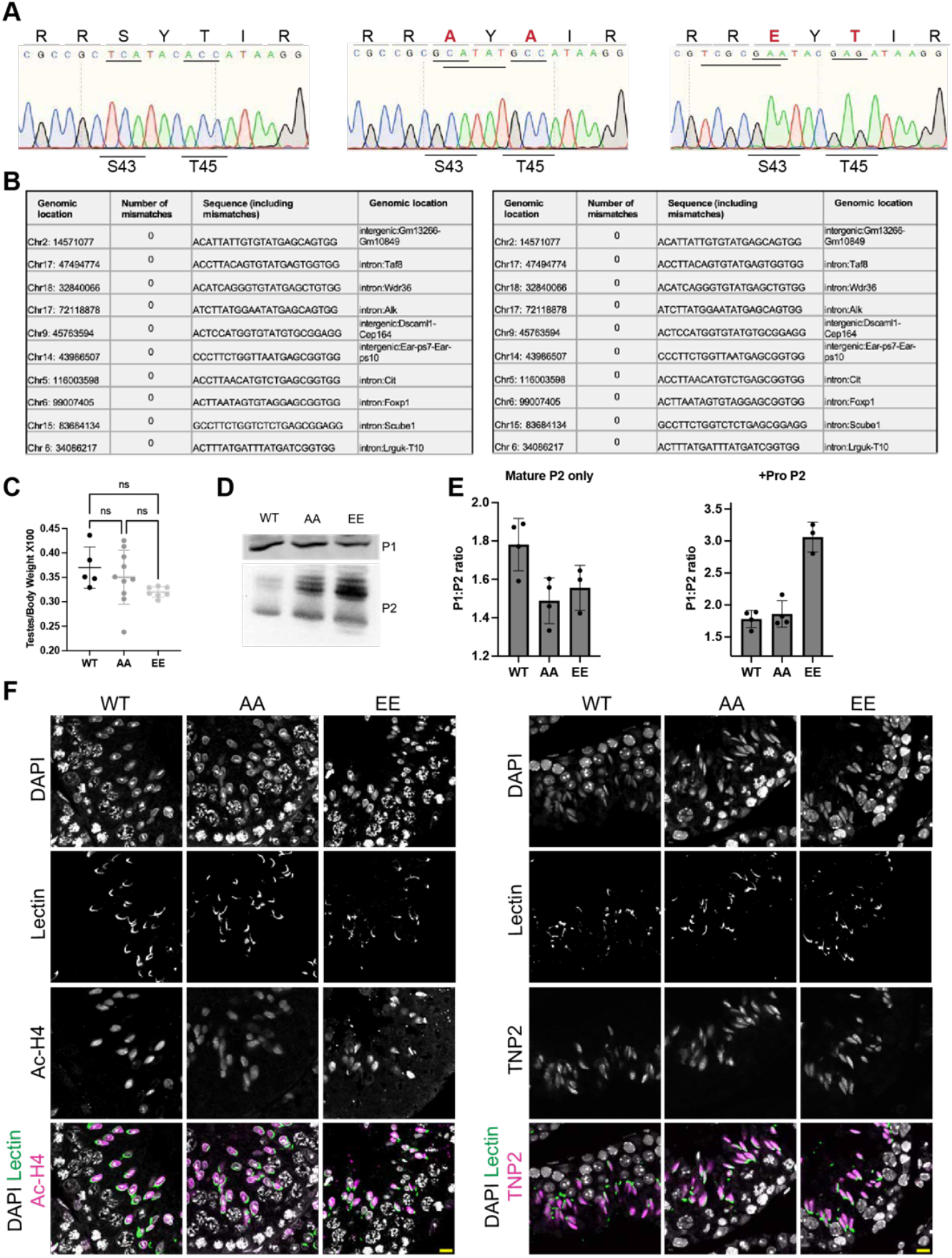
P1 S43/T45 substitutions do not affect P1 abundance, testis size, or initiation of histone-to-protamine exchange. **(A)** Representative Sanger sequencing traces showing successful substitution of S43 and T45 to either alanine or glutamic acid. **(B)** Sequencing of the top 10 potential off-target sites from both AA (left) and EE (right) males shows no off-target genetic modifications present. **(C)** Testis/body weight ratio from WT (n=5), AA (n=7), and EE (n=5) males. Each dot represents the measurement of a single animal. Statistical tests were performed using ANOVA and adjusted for multiple comparisons. Center line represents the mean and error bars represent standard deviation. **(D)** Immunoblot of protein lysates from adult WT, AA, and EE testes shows no defects in P1 expression between genotypes. **(E)** Quantification of P1:P2 ratio from WT, AA, and EE sperm. Left plot depicts P1: mature P2 ratio (not including residual pro P2) and right plot depicts a corrected P1:P2 ratio including residual pro P2. **(F)** Immunofluorescence of WT, AA, and EE testes cross sections stained for Ac-H4 (left) or TNP2 (right) highlights normal progression of histone-to-protamine exchange. Scale bar: 10 μm

**Supplemental Figure 5:**
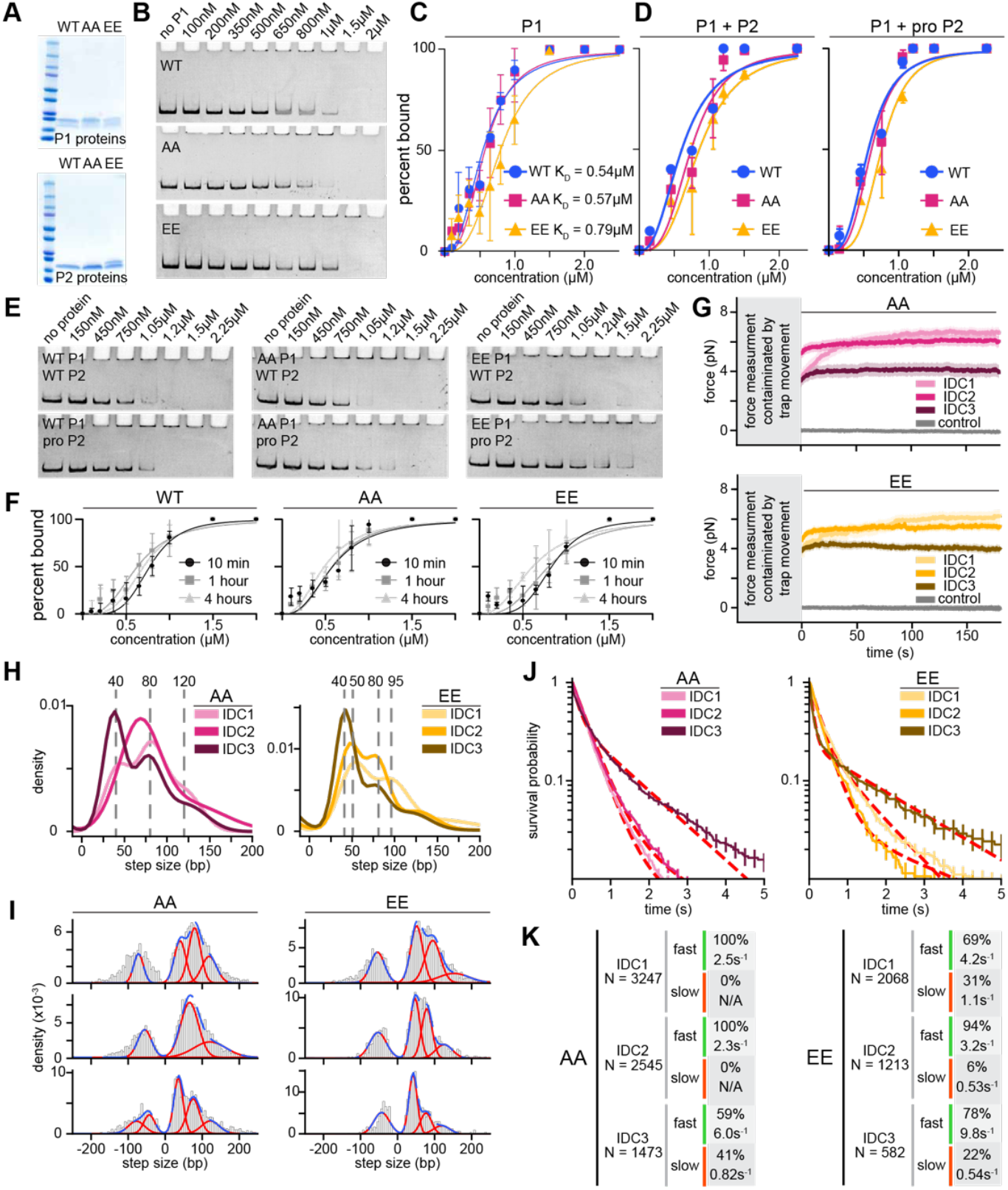
P1 S43/T45 charge states fine-tune DNA-binding strength and chromatin compaction/decompaction dynamics. **(A)** Coomassie-stained gel of purified P1 (top) and P2 (bottom) proteins purified from WT, AA, and EE sperm. **(B)** Representative EMSAs of increasing amounts of WT P1 (top), AA (middle), and EE (bottom) a with a ≈300 bp DNA fragment. **(C)** Quantification of the binding affinities of WT P1, AA, and EE to a ≈300 bp DNA fragment. K_D,app_ values were calculated using the Hill equation. **(D)** Quantification of the binding affinities of WT and mutant P1 mixed at a 1:2 ratio with P2 derived from either WT or EE sperm. **(E)** Representative EMSAs of WT and each mutant P1 mixed at a 1:2 ratio with P2 derived from WT (top) or EE (bottom) sperm at increasing total protamine concentration. **(F)** Quantification of DNA binding EMSAs of WT P1, AA, and EE after 10 minutes, 1 hour, or 4 hours of incubation illustrating that protamine-DNA complexes reach an equilibrium within 10 minutes. **(G)** Force creep curves for AA (top) and EE (bottom) measured during the incubation phase of all three IDCs. **(H)** Joint histogram of AA (left) and EE (right) decompaction steps from all three IDCs, grey dashed lines indicate peaks from Gaussian fits. **(I)** Complete histograms of AA (left) and EE (right) decompaction steps from each IDC. Individual Gaussian fits presented in red and combined fit in blue. **(J)** Survival probability of step dwell times for AA (left) and EE (right) decompaction steps from all IDCs, red dashed lines indicate bi-exponential fits. **(K)** Tables of kinetic parameters for AA (left) and EE (right) from each IDC.

**Supplemental Figure 6:**
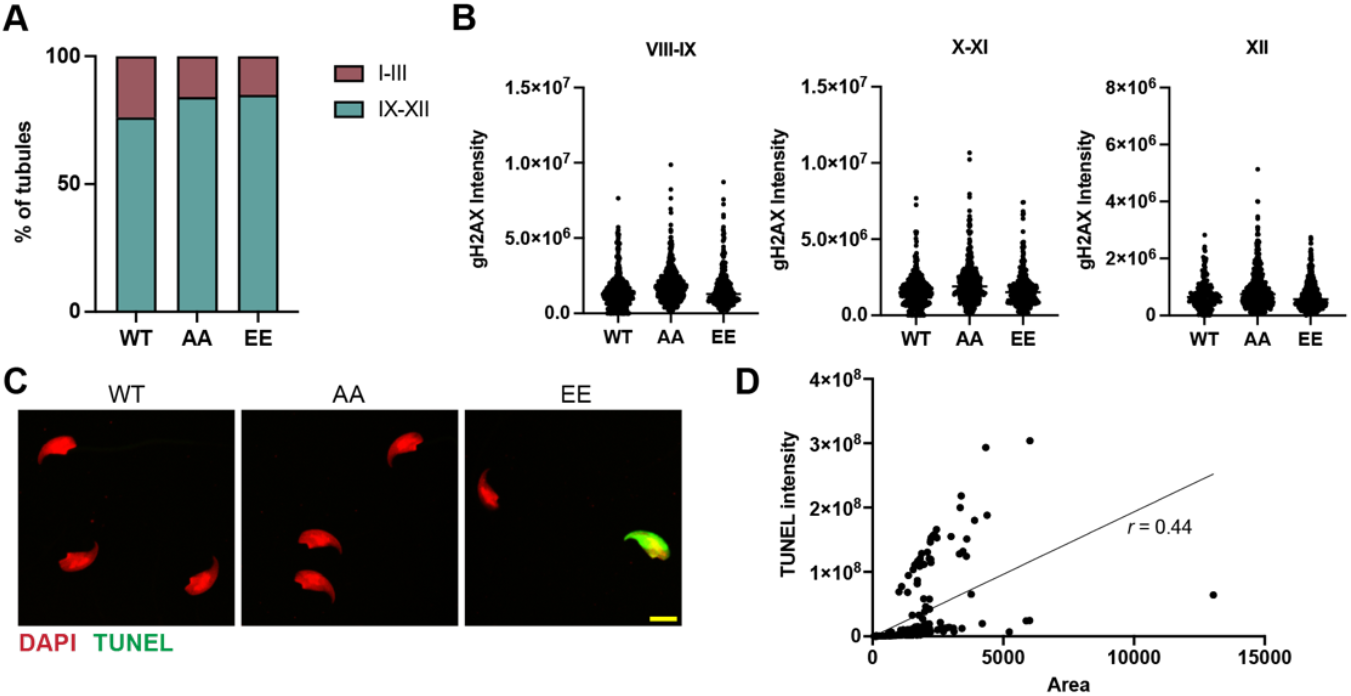
P1 mutants repair programmed DNA breaks during spermiogenesis, but EE mutants accrue damage outside the testis. **(A)** Quantification of seminiferous tubule stages observed in synchronized males used for DNA damage assessment by Comet in Figure 6C. **(B)** Quantification of gH2AX levels in different stages of testicular spermatids highlights a comparable level of programmed damage in stages 9-11 that are largely resolved by stage 12 across all genotypes. **(C)** Representative immunofluorescence of TUNEL+ EE sperm as a marker of DNA damage. Scale bar: 5 μm. **(D)** Correlation of TUNEL staining intensity and overall sperm area of EE sperm shows a positive correlation, r=0.44.

## Notes

### Competing Interest Statement

The authors have declared no competing interest.

